# Functional connectivity of the inferior frontal gyrus: A meta-analytic connectivity modeling study

**DOI:** 10.1101/2022.02.17.480832

**Authors:** Talat Bulut

## Abstract

**Background:** Neurocognitive models of language processing highlight the role of the left inferior frontal gyrus (IFG) in the functional network underlying language. Furthermore, neuroscience research has shown that IFG is not a uniform region anatomically, cytoarchitectonically or functionally. However, no previous study explored the language-related functional connectivity patterns of different IFG subdivisions using a meta-analytic connectivity approach.

**Purpose:** The present meta-analytic connectivity modeling (MACM) study aimed to identify language-related coactivation patterns of the left and right IFG subdivisions.

**Method:** Six regions of interest (ROIs) were defined using a probabilistic brain atlas corresponding to pars opercularis (BA44), pars triangularis (BA45) and pars orbitalis (Fo6&7) of IFG in both hemispheres. The ROIs were used to search the BrainMap functional database to identify neuroimaging experiments with healthy, right-handed participants reporting language-related activations in each ROI. Activation likelihood estimation analyses were then performed on the foci extracted from the identified studies to compute functional convergence for each ROI, which was also contrasted with the other ROIs within the same hemisphere. In addition, a behavioral analysis was conducted to determine functional specificity for language subdomains within each ROI.

**Results:** A primarily left-lateralized functional network was revealed for the left and right IFG subdivisions. The left IFG ROIs exhibited a more robust coactivation pattern than the right IFG ROIs. In particular, the left posterior-dorsal IFG (BA44) was associated with the most extensive coactivation pattern involving bilateral frontal, bilateral parietal, left temporal, left subcortical (thalamus and putamen), and right cerebellar regions, while the left anterior-ventral IFG (BA45 and Fo6&7) revealed a predominantly left-lateralized involvement of frontotemporal regions.

**Conclusion:** The findings align with the neurocognitive models of language processing that propose a division of labor among the left IFG subdivisions and their respective functional networks. Also, the opercular part of left IFG (BA44) stands out as a major hub in the language network with connections to diverse cortical, subcortical and cerebellar structures.

## Introduction

### Left inferior frontal gyrus and its role in language processing

It is generally assumed that Broca’s area consists of pars opercularis (BA44) and pars triangularis (BA45) of the left inferior frontal gyrus (LIFG) (Amunts et al., 1999). However, due to its involvement in language processing, pars orbitalis (BA47) of LIFG has also been viewed as part of a broadly-defined Broca’s complex (Xiang, Fonteijn, Norris, & Hagoort, 2010). Previous neuroimaging studies associated parts of this region with syntactic processing (Dapretto & Bookheimer, 1999; Pallier, Devauchelle, & Dehaene, 2011; Segaert, Menenti, Weber, Petersson, & Hagoort, 2012), semantic processing (Dapretto & Bookheimer, 1999; Zhang et al., 2019; Zhu et al., 2019), phonological processing (Heim, Opitz, Müller, & Friederici, 2003; Matsuo et al., 2010), and morphological processing (Bozic, Fonteneau, Su, & Marslen-Wilson, 2015; Laine, Rinne, Krause, Teräs, & Sipilä, 1999; Sahin, Pinker, & Halgren, 2006), among others. In particular, LIFG has frequently been associated with syntactic processing. In support of this association, previous meta-analyses and systematic reviews of neuroimaging research on syntax revealed functional convergence in LIFG, in addition to several other regions including the posterior temporal lobe (Grodzinsky, Pieperhoff, & Thompson, 2021; Hagoort & Indefrey, 2014; Meyer & Friederici, 2016; Rodd, Vitello, Woollams, & Adank, 2015; Walenski, Europa, Caplan, & Thompson, 2019; Zaccarella, Schell, & Friederici, 2017).

Previous research has shown that LIFG is not a uniform region anatomically, cytoarchitectonically or functionally (Amunts et al., 1999; Clos, Amunts, Laird, Fox, & Eickhoff, 2013; Wojtasik et al., 2020). Along similar lines, several studies found that different LIFG subdivisions responded to different linguistic processes, usually involving posterior-dorsal LIFG (BA44) with syntactic processing and anterior-inferior LIFG (BA45 and/or BA47) with semantic processing (Dapretto & Bookheimer, 1999; Schell, Zaccarella, & Friederici, 2017). Some of the previous meta-analyses cited in the preceding paragraph also support functional differentiation within LIFG (Hagoort & Indefrey, 2014; Zaccarella et al., 2017; c.f. Rodd et al., 2015), since convergence for semantically challenging sentences was located more anteriorly (BA45) than that for syntactically challenging sentences (BA44) (Hagoort & Indefrey, 2014), and since foci activated for the comparison of syntactically licit sequences vs. word lists converged more strongly in BA44 (Zaccarella et al., 2017). Similarly, an activation likelihood estimation meta-analysis of neuroimaging experiments on inflectional morphology also revealed more robust involvement of LBA44 in morphological processing than other LIFG subdivisions, supporting the correlation between posterior LIFG and grammatical processing (Bulut, in review). The functional subdivision of LIFG into ventral-dorsal or anterior-inferior-posterior portions has been taken up by several neurocognitive accounts of language processing, as addressed in the following section.

### Neurocognitive models of language processing

Several models have been conceived to explain the neural circuitry of language processing. In all the models reviewed in this section, LIFG plays a pivotal role with functions attributed to this region ranging from syntactic and semantic processing (Friederici, 2002, 2011, 2012), unification of phonological, syntactic and semantic information (Hagoort, 2013, 2016; Xiang et al., 2010), to speech production (Hickok & Poeppel, 2004, 2007; Poeppel, Emmorey, Hickok, & Pylkkanen, 2012), and processing declarative (lexical) and procedural (grammatical) information (Ullman, 2001, 2004, 2016). However, all these models stress that these functions arise through connectivity of LIFG with certain other regions as part of a functional network, as detailed below.

The neurocognitive model of language processing proposed by Friederici posits that during auditory sentence comprehension, linguistic information received by the primary auditory cortex flows through two distinct ventral and dorsal pathways in the left hemisphere (Friederici, 2002, 2011, 2012). Ventrally, the anterior superior temporal gyrus (STG), the middle temporal gyrus (MTG) and anterior-ventral LIFG (BA45 and BA47) underlie semantic processing, while posterior-dorsal LIFG (BA44) and posterior STG / superior temporal sulcus (STS) constitute the dorsal pathway responsible for syntactic and grammatical processing. Recent versions of this account also conceptualize a second dorsal pathway which connects posterior STG with premotor cortex directly, and indirectly via inferior parietal cortex (Friederici, 2011, 2012). The premotor cortex is further connected with LBA44. It is claimed that this second dorsal pathway facilitates mapping between phonological and motor/articulatory information. Thus, Friederici’s model predicts that anterior-ventral LIFG (BA45/47) will coactivate with anterior STG and MTG since they comprise the ventral pathway, while posterior-dorsal LIFG (BA44) will exhibit functional connectivity with posterior STG/STS as part of the first dorsal pathway, and with the premotor and inferior parietal cortices, as part of the second dorsal pathway.

The Memory, Unification and Control (MUC) Model put forward by Hagoort, on the other hand, associates the left temporal lobe with memory processes and LIFG with unification processes underlying language (Hagoort, 2005, 2013, 2016). According to this view, the memory areas within the left temporal lobe embodies the mental lexicon by storing lexical information, while the unification areas within LIFG is responsible for combinatorial processing of linguistic information. Executive control processes such as attention, selection, turn-taking in conversation and code-switching underlying language are subserved by the control network spanning the dorsolateral prefrontal cortex; i.e., the middle frontal gyrus (MFG) (BA46, 9, 8, 6), the anterior cingulate cortex and parts of the parietal cortex (Hagoort, 2005, 2013, 2016; Xiang et al., 2010). The MUC Model emphasizes dynamic interaction between the memory, unification and control processes. Importantly, gradients within the frontotemporal regions are proposed that map phonological processes dorsally, semantic processes ventrally and syntactic processes in between these ventral and dorsal sites (Hagoort, 2013, 2016; Xiang et al., 2010). Specifically, the model associates LBA44 and the left posterior STG/STS with phonological processing, LBA45 and the left posterior MTG with syntactic processing, and LBA47 and the left posterior inferior temporal gyrus (ITG) with semantic processing. Extensions of the model also suggest similar ventral to dorsal gradients within the parietal lobe extending from the angular gyrus to the supramarginal gyrus, and within the dorsolateral prefrontal cortex extending from anterior-inferior MFG to posterior-superior MFG (Hagoort, 2013, 2016; Xiang et al., 2010), which exhibit functional connectivity with the corresponding ventral-dorsal LIFG subdivisions for semantic, syntactic and phonological processes. Therefore, the MUC Model predicts that in the left hemisphere, posterior-dorsal IFG (LBA44) will coactivate with dorsal temporal (posterior STG/STS), dorsal parietal (supramarginal gyrus) and posterior dorsolateral prefrontal (posterior-superior MFG) regions. The model also predicts that ventral LIFG (BA47) will coactivate with ventral temporal (posterior ITG), ventral parietal (angular gyrus) and anterior dorsolateral prefrontal (anterior-inferior MFG) regions. Finally, anterior LIFG (BA45) is predicted to coactivate with the temporal, parietal and dorsolateral prefrontal regions intermediate with the respective ventral-dorsal aspects.

Another account that emphasizes a distinction between the lexicon and combinatorial processing is the Declarative/Procedural (DP) Model put forward by Ullman (Ullman, 2001, 2004, 2016). The DP Model distinguishes between a declarative system, i.e., the mental lexicon, responsible for learning, consolidating and storing lexical information, and a procedural system, i.e., grammar, responsible for rule-governed grammatical processing. The model incorporates a declarative system in which medial temporal lobe structures including the hippocampus are responsible for learning and consolidating linguistic information, which is then stored in the temporal cortex laterally. The declarative system also implicates anterior-inferior portions of IFG (BA45/47) in encoding, selection and retrieval of declarative memories (Ullman, 2004). According to the DP Model, the procedural system underlying rule-governed grammatical processing depends on a frontal-basal ganglia circuit. The procedural system is subserved by basal ganglia structures (caudate nucleus and anterior putamen) and, to a certain extent, by the cerebellum, which are primarily involved in learning and consolidation of procedural memories, as well as by the left premotor cortex (BA6) and posterior LIFG (BA44), which are claimed to process automatized procedural memories (Ullman, 2016). Thus, the DP Model predicts that anterior-inferior IFG (BA45/47) will coactivate with the lateral and medial temporal lobe as part of the declarative system, while posterior LIFG (BA44) will coactivate with the left premotor cortex, the basal ganglia and the cerebellum.

Another account linking linguistic function to specific brain regions and their connections is the Dual-Stream Model of Speech Processing (Hickok & Poeppel, 2004, 2007; Poeppel et al., 2012). According to this account, auditory input registered by the primary auditory cortex is subjected to phonological analysis at the middle to posterior portions of superior STS bilaterally. Following phonological processing, the information flows through two distinct systems, a largely bilateral ventral system underlying speech comprehension (mapping phonological information onto meaning and concepts), and a left-lateralized dorsal stream underlying speech production (mapping phonological information to articulatory motor representations). The ventral stream involves bilateral temporal regions, with the posterior middle and inferior temporal cortices associated with phonological-semantic mappings, and the left anterior temporal lobe associated with combinatorial processing of syntactic and semantic information. The dorsal stream, however, spans area Spt (a region of the Sylvian fissure at the parieto-temporal boundary) responsible for sensorimotor interface as well as dorsal LIFG (BA44 and 45), the left premotor cortex and the left insula forming the articulatory network (Hickok & Poeppel, 2007; Poeppel et al., 2012). According to this model, there is a direct connection between the ventral and dorsal streams via the left anterior temporal lobe and dorsal LIFG. Hence, the Dual-Stream Model predicts that dorsal LIFG (BA44 and 45) will coactivate with area Spt, the left premotor cortex and the left insula as part of the dorsal stream, and also with the left anterior temporal lobe, with which it has a direct connection.

In summary, all neurocognitive models of language processing reviewed above suggest a division of labor within LIFG. That is, anterior-ventral LIFG (usually BA47, but sometimes BA45, as well) is associated with several functions including semantic processing (Friederici, 2011, 2012; Hagoort, 2013), processing declarative memories involving lexical information (Ullman, 2001, 2004, 2016), while posterior-dorsal LIFG (usually BA44, but sometimes BA45, as well) is claimed to underlie syntactic and grammatical processing (Friederici, 2002, 2011, 2012; Hagoort, 2005, 2013, 2016; Ullman, 2001, 2004, 2016), phonological processing (Hagoort, 2005, 2013, 2016), and speech production (Hickok & Poeppel, 2004, 2007; Poeppel et al., 2012). The models also exhibit a certain degree of similarity in their claims of ventral versus dorsal connectivity patterns for the posterior-dorsal versus anterior-ventral portions of LIFG. Despite these similarities, however, there are substantial differences among the models regarding the regions included in these ventral and dorsal networks across the temporal, parietal and frontal lobes, as well as regarding the involvement of subcortical structures, as summarized above. Finally, the language network proposed by the accounts presented here is strongly left-lateralized, with only limited involvement of the right hemisphere (Hickok & Poeppel, 2004, 2007; Poeppel et al., 2012).

### Functional and structural connectivity of LIFG

Although LIFG has been central to debates about the relationship between language and the brain, the neurocognitive models of language processing presented above highlight the importance of a functional network for language. Indeed, modern neuroscience has moved away from attributing certain functions to isolated brain regions towards appreciating functional circuitry across the brain underlying cognitive functions (Viñas-Guasch & Wu, 2017). Accordingly, the function of LIFG should also be conceptualized as part of a functional and structural neural network for language (Friederici, 2011).

Several techniques have been developed and frequently used to identify structural and functional connectivity patterns in the brain. Structural connections among brain regions have been investigated in vivo using diffusion tensor imaging (DTI), which involves a special use of MRI to identify white matter fibers by measuring diffusion of water molecules (Alexander, Lee, Lazar, & Field, 2007; van den Heuvel & Hulshoff Pol, 2010). Functional connectivity, on the other hand, refers to functional communication or correlated neural oscillations between anatomically distinct brain regions, and has usually been studied using resting-state fMRI (r-s fMRI), which estimates spontaneous low-frequency fluctuations in the BOLD signal (Lee, Smyser, & Shimony, 2013; van den Heuvel & Hulshoff Pol, 2010). It is generally assumed that functional connections strongly correlate with structural white matter connections (Greicius, Supekar, Menon, & Dougherty, 2009; Huang & Ding, 2016; van den Heuvel & Hulshoff Pol, 2010).

A substantial number of DTI studies investigated the structural connectivity underlying language (for reviews, please see Axer, Klingner, & Prescher, 2013; Dick & Tremblay, 2012). This body of research confirmed the classical language pathway between Broca’s area and Wernicke’s area via arcuate fasciculus as part of the superior longitudinal fasciculus, which was found to be stronger in the left hemisphere (Glasser & Rilling, 2008; Parker et al., 2005; Powell et al., 2006). Furthermore, it has been shown that distinct dorsal and ventral fiber tracts connect IFG with temporal (usually superior and middle temporal gyri) and inferior parietal (supramarginal and angular gyri) cortices, dorsal connections mediated by arcuate / superior longitudinal fasciculi, and ventral connections by several fiber tracts including extreme capsule, external capsule, uncinate fasciculus, inferior fronto-occipital fasciculus, and middle longitudinal fasciculus (Axer et al., 2013; Kellmeyer et al., 2013; Parker et al., 2005; Saur et al., 2010). Moreover, it has also been reported that the dorsal pathway originates within posterior IFG (BA44), while the ventral pathway originates within anterior IFG (BA45) and sometimes also within inferior IFG (BA47) (Kellmeyer et al., 2013; Saur et al., 2010). These dorsal versus ventral connections have been interpreted as underlying phonological-articulatory processes and lexical-semantic processes, respectively (Kellmeyer et al., 2013; Saur et al., 2010). However, there is still controversy regarding interpretation of the functional roles that these fiber bundles play (Axer et al., 2013).

Investigations of structural connections among brain regions have been complemented by explorations of functional connectivity, which identified distinct functional networks such as motor, visual, auditory, default mode and attention networks (van den Heuvel & Hulshoff Pol, 2010). Several studies also revealed a functionally connected, mostly perisylvian network using r-s fMRI (Tomasi & Volkow, 2012; Xiang et al., 2010). In parallel with the structural network, these studies identified a mostly left-lateralized functional network associated with Broca’s and Wernicke’s areas (Tomasi & Volkow, 2012; Xiang et al., 2010). Specifically, it was shown that the activity of Broca’s and Wernicke’s areas correlated with the inferior and middle frontal gyri, the inferior and superior temporal cortices, the inferior parietal cortex as well as subcortical regions including the basal ganglia and the cerebellum (Tomasi & Volkow, 2012). Another r-s fMRI study examined the functional connectivity patterns of six seed regions: pars opercularis, triangularis and orbitalis of IFG in both hemispheres (Xiang et al., 2010). This study found that IFG coactivated with a widespread network spanning frontal (IFG, MFG, and precentral gyrus), insular, posterior temporal (STG, MTG, and ITG), and parietal (superior and inferior parietal lobules) cortices as well as subcortical regions (caudate and putamen). Importantly, a dorsal-to-ventral functional connectivity pattern was observed for distinct IFG subdivisions in the posterior temporal (posterior STG, posterior MTG, and posterior ITG), superior and inferior parietal (postcentral, supramarginal and angular gyri) and dorsolateral prefrontal (MFG) cortices. It was claimed that these dorsal-ventral gradients of functional connectivity for the posterior, anterior and inferior parts of IFG were stronger in the left hemisphere and corresponded to phonological, syntactic and semantic processes, respectively, in support of the MUC Model described above (Hagoort, 2005, 2013, 2016).

To summarize, investigations of structural and functional connectivity provided fairly consistent evidence for dorsal and ventral connectivity patterns correlated with LIFG. However, several limitations prevent delineating the specifics of a language network based on these studies. For instance, the limited resolution of DTI may pose challenges to linking white matter fibers to specific grey matter sites (Ford et al., 2013; Xiang et al., 2010). Furthermore, the extent to which functional and structural connections revealed in these studies underlie language is questionable as DTI and r-s fMRI studies identify task-independent general connectivity patterns that may or may not apply to certain cognitive domains. In particular, the inferior frontal cortex and its connections have been associated not only with language, but also with domain-general functions ranging from attention, action processing, working memory, cognitive control to processing emotions, music, math and numbers (Belyk, Brown, Lim, & Kotz, 2017; Clos et al., 2013; Hung et al., 2015; Klimovich-Gray & Bozic, 2019; Maess, Koelsch, Gunter, & Friederici, 2001; Maruyama, Pallier, Jobert, Sigman, & Dehaene, 2012; Novick, Trueswell, & Thompson-Schill, 2005, 2010; Papitto, Friederici, & Zaccarella, 2020; Sebastian et al., 2016; Sundermann & Pfleiderer, 2012; Thothathiri, Schwartz, & Thompson-Schill, 2010). Therefore, IFG arguably assumes its domain-specific linguistic function as a constituent of a wider domain-specific network (Friederici, 2011).

### Meta-analytic connectivity modeling and the current study

A recent development in neuroimaging research is meta-analytic connectivity modeling (MACM), which has been increasingly utilized to explore functional connectivity of a given region of interest. MACM is an extension of the activation likelihood estimation (ALE) method frequently used in meta-analyses of neuroimaging experiments on a particular cognitive function or task. Coordinate-based meta-analyses such as ALE and MACM facilitate integration of neuroimaging experiments to achieve findings with high generalizability (Samartsidis et al., 2020). In the conventional ALE method, a meta-analysis of previously published neuroimaging studies is conducted to identify functional convergence in the brain across those studies for a given task or cognitive process (Laird, Fox, et al., 2005; Müller et al., 2018; Turkeltaub, Eden, Jones, & Zeffiro, 2002). In the MACM approach, however, functional convergence of studies that report activation in a predetermined region of interest (ROI) is computed (Robinson, Laird, Glahn, Lovallo, & Fox, 2010). Hence, MACM allows identification of coactivation patterns, or functional connectivity, of an ROI, usually independently of a task or cognitive process. This method has been used to explore task-independent functional connectivity of several target regions (Erickson, Rauschecker, & Turkeltaub, 2017; Robinson et al., 2010). More recently, MACM has been employed to identify coactivation of a given brain region in a given cognitive domain such as emotion (Belyk et al., 2017) and language (Ardila, Bernal, & Rosselli, 2016; Bernal, Ardila, & Rosselli, 2015; Viñas-Guasch & Wu, 2017). For instance, an MACM study of LBA44 revealed mainly left-lateralized functional connectivity across frontal, inferior and superior parietal, and posterior temporal regions (Bernal et al., 2015). Other ALE and MACM studies reported functionally distinct clusters within LBA44 and suggested further functional segregation of LBA44 into anterior and posterior regions, responsible for language and action, respectively, and each associated with differential functional networks (Clos et al., 2013; Papitto et al., 2020). A similar subdivision was also suggested for pars orbitalis (BA47) of LIFG in another study combining Kernel Density Estimation meta-analysis with MACM (Belyk et al., 2017). This study associated lateral LBA47 with both semantic and emotional processing, and opercular LBA47 with emotional processing alone. A functional connectivity network largely shared between these two subdivisions was reported, though the lateral zone exhibited more pronounced prefrontal connectivity.

Despite these advances in our understanding of the language network and the role of IFG in it, there is still much that remains to be explored. In particular, limited research explored meta-analytic connectivity of LIFG in language tasks, while no meta-analysis was conducted on language-related functional connectivity of the right inferior frontal gyrus (RIFG). Furthermore, functional connectivity patterns of different IFG subdivisions (opercular, triangular and orbital parts) have not been addressed using a meta-analytic connectivity approach. To fill these gaps, the present MACM study aimed to identify language-related functional connectivity or coactivation patterns of the left and right IFG subdivisions in healthy participants. To that end, six regions of interest (ROIs) were defined using a probabilistic brain atlas corresponding to pars opercularis (BA44), pars triangularis (BA45) and pars orbitalis (BA47) of IFG in both hemispheres. The ROIs and several criteria were used to search the BrainMap functional database to identify experiments reporting language-related activations in each ROI. ALE analyses were then performed on the foci extracted from the identified studies to compute functional convergence for each one of the ROIs, which was then contrasted with the other ROIs within the same hemisphere. Exploration of functional connectivity of IFG subdivisions across language tasks can shed light on the neural network underlying language and constrain neurocognitive models of language processing. In addition, delineating the functional network of language processing in health may provide a baseline against which to compare the language network in clinical populations and assess neurorehabilitation in disorders of language such as aphasia.

## Materials and methods

The current study used the MACM method to examine the functional connectivity of three subdivisions of IFG in each hemisphere. For this purpose, the following steps were carried out: Six ROIs (three in each hemisphere) were defined corresponding to opercular, triangular and orbital aspects of IFG based on a cytoarchitectonic, probabilistic brain atlas. The predefined ROIs and additional search criteria were entered in Sleuth to search the BrainMap database in order to identify experiments reporting language-related activations that include the relevant ROI (as well as activations elsewhere in the brain). The activation foci in the identified experiments, which included activations not only within the ROI but also elsewhere in the brain, were extracted using Sleuth and entered in GingerALE, where ALE analyses were performed to identify coactivation network of each ROI. Afterwards, the coactivation network of each ROI was contrasted with the other ROIs within the same hemisphere.

### Regions of interest

Six regions of interest (ROIs) were defined using the probabilistic Julich-Brain atlas, which is based on the brain’s cytoarchitecture (Amunts, Mohlberg, Bludau, & Zilles, 2020). Probabilistically defined ROIs offer several advantages over functionally or geometrically defined ones, as the former enable a more consistent characterization across studies and ensure better localization of the target structures (Robinson et al., 2010). In the present study, the ROIs comprised three structures within LIFG and RIFG, each (Figure 1). LBA44 and RBA44 comprised the posterior portion of IFG, i.e., pars opercularis, in the left and the right hemispheres, respectively. LBA45 and RBA45 constituted the anterior portion of IFG, i.e., pars triangularis, in the left and the right hemispheres, respectively. More ventrally, LFo6&7 and RFo6&7 spanned parts of the lateral orbitofrontal cortex primarily including BA47, i.e., pars opercularis (Wojtasik et al., 2020). The ROI maps in the MNI Colin 27 template were downloaded from the European Human Brain Project website (https://ebrains.eu), where cytoarchitectonic brain regions are curated and constantly updated (Amunts et al., 2020). The latest versions of the maps as of the preparation of the present study were used; i.e., v9.2 for the left and right BA44 and BA45, and v3.2 for the left and right Fo6&7. The maps were then overlaid on the MNI template (Colin27_T1_seg_MNI.nii) available on GingerALE’s website using the Mango software (http://ric.uthscsa.edu/mango/) (Lancaster et al., 2010). LFo6&7 and RFo6&7 were created by combining the Fo6 and Fo7 maps downloaded from the European Human Brain Project website. The probabilistic maps were then thresholded using different probabilities for each ROI (see the color bars in Figure 1) such that each region had a probability greater than 0.48, with mean probabilities ranging from 0.64 to 0.77, and a similar size^1^ ranging from 2310mm^3^ to 2409mm^3^. Therefore, a balance was struck between the volumetric sizes of the ROIs and their probability of representing respective brain regions to enable direct comparisons among the ROIs. Importantly, thresholding also ensured that there was no overlap among the ROIs. The ROIs were then created based on the thresholded maps. Finally, the ROIs were visually inspected using the Talairach Daemon in Mango (Lancaster et al., 1997; Lancaster et al., 2000) and using different brain templates in MRIcron (https://www.nitrc.org/projects/mricron) (Rorden & Brett, 2000) to ensure that the intended brain regions were captured.

**Figure 1.**
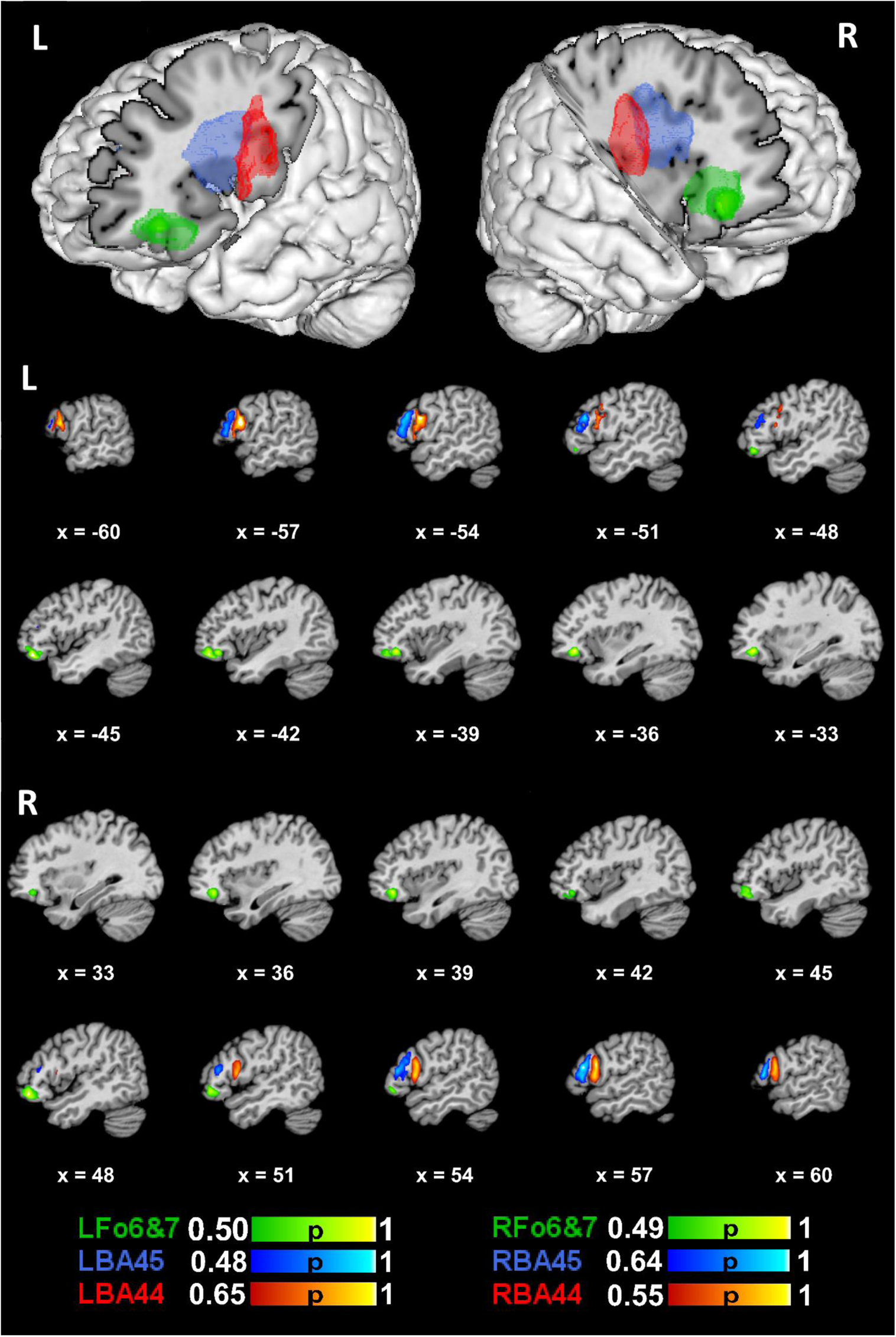
Anatomical 3-D renderings of the left and right hemispheric ROIs used in the meta-analysis. BA44 is represented in red, BA45 in blue and Fo6&7 in green for both left and right hemispheres. The color bars indicate probability of the relevant structure to be found in that area. The figure was produced using the Mango software (http://ric.uthscsa.edu/mango/).

### Database search

A search was conducted within the BrainMap functional database on 9-27-2021 using Sleuth Version 3.0.4 (Fox et al., 2005; Fox & Lancaster, 2002; Laird, Lancaster, & Fox, 2005). At the time of the search, the functional database comprised 3406 papers, 16901 experiments, 76016 subjects and 131598 locations. The BrainMap taxonomy allows viewing and automatically searching meta-data of articles submitted to the database including imaging modality (e.g., PET or fMRI), subjects (e.g., handedness, age, gender), and behavioral domains (e.g., action, cognition, emotion) as well as behavioral subdomains (e.g., cognition-attention, cognition-language, cognition-language-semantics) (Fox et al., 2005; Lancaster et al., 2012). The database search was intended to identify studies on language recruiting only right-handed subjects to identify language-specific connectivity patterns of the ROIs. Thus, the following search query was used: “locations: left and right IFG ROIs”, “experimental context: normal mapping”, ‘‘behavioral domain: cognition-language”, “experimental activation: activations only”, “subjects: normals”, “handedness: right”, “imaging modality: fMRI or PET”. Restriction of the search to “normals” ensured that only the experiments conducted with healthy subjects were included. The ROIs defined as explained above were separately included as a search criterion in the database searches. The search intended to yield only language-specific activations; hence, only the “cognition-language” behavioral domain encompassing all linguistic levels (phonology, orthography, semantics, syntax, speech) was used, but not “action-execution-speech”, which would mainly involve action-related processes of articulation. The results identified by the database search are summarized in Table 1 below.

**Table 1.** Database search results for each ROI.

| ROI | Papers | Subjects | Experiments | Locations |
| --- | --- | --- | --- | --- |
| LBA44 | 72 | 1100 | 95 | 1394 |
| LBA45 | 68 | 926 | 88 | 1264 |
| LFo6&7 | 29 | 436 | 34 | 538 |
| RBA44 | 22 | 279 | 30 | 492 |
| RBA45 | 21 | 298 | 23 | 333 |
| RFo6&7 | 14 | 240 | 14 | 221 |

The distribution across behavioral subdomains of language-related experiments identified for each ROI and entered in the MACM analyses is shown in Table 2. Please note that an experiment can pertain to more than one language subdomain (e.g., both phonology and semantics), depending on the specifics of the contrast and stimuli used. Table 2 also reports the distribution of experiments across language subdomains in the entire BrainMap database, which was identified using the same search criteria (e.g., handedness) as those for the ROIs except that the ROIs were removed from the search query to identify all language-related experiments in the database reporting coordinates anywhere in the brain. Experiments which were not assigned to a language subdomain, but were solely assigned to the general domain of language are indicated in the ‘unspecified’ row in Table 2. The Sleuth workspace files with metadata of the experiments included in each ALE analysis are accessible at https://www.doi.org/10.17632/nyx2gz9yww.1.

**Table 2.** Distribution across language subdomains of the experiments entered in the MACM analysis for each ROI and overall distributions across the BrainMap database.

| Language Subdomain | LBA44 | LBA45 | LFo6&7 | RBA44 | RBA45 | RFo6&7 | BrainMap |
| --- | --- | --- | --- | --- | --- | --- | --- |
| Orthography | 10 | 8 | 4 | 2 | 2 | 2 | 188 |
| Phonology | 25 | 8 | 4 | 7 | 3 | 2 | 295 |
| Semantics | 37 | 59 | 27 | 17 | 15 | 11 | 908 |
| Speech | 38 | 30 | 5 | 11 | 7 | 2 | 627 |
| Syntax | 10 | 5 | 2 | 1 | 3 | 1 | 112 |
| Unspecified | 6 | 6 | 0 | 1 | 1 | 0 | 59 |

The coordinates identified in each search were grouped by experiment (Turkeltaub et al., 2012) and exported as a text file to be input in GingerALE. The icbm2tal transform was used to automatically convert coordinates reported in Talairach space into MNI space (Laird et al., 2010; Lancaster et al., 2007).

### ALE analyses

GingerALE 3.0.2 was used (Eickhoff, Bzdok, Laird, Kurth, & Fox, 2012; Eickhoff et al., 2009) to estimate the convergence of coactivations for each ROI. Therefore, six separate ALE analyses were conducted with the activation coordinates obtained for the ROIs. The ALE analyses were performed according to the standard procedures reported in previous research (Cieslik, Mueller, Eickhoff, Langner, & Eickhoff, 2015; Müller et al., 2017; Wojtasik et al., 2020). Thus, 3D Gaussian probability distributions centered at each group of foci were created with a full-width half-maximum (FWHM) that was computed based on the sample size in each experiment (Eickhoff et al., 2009). Next, voxel-wise ALE scores were calculated by obtaining the union of modeled activation maps for each experiment. The union of these activation probabilities were then compared against the null hypothesis of a random spatial relationship between the experiments. The *p*-value distributions resulting from these probabilities were then thresholded at a voxel-level uncorrected cluster-forming threshold of *p* < 0.001 and a cluster-level corrected threshold of *p* < 0.05 (family-wise error-corrected for multiple comparisons), with 10000 thresholding permutations.

Following the coactivation analyses separately for each ROI, the conjunction/intersection of coactivations within each hemisphere was visualized using the “Overlay Logicals” utility of the Mango software (Lancaster et al., 2010). Specifically, the intersection of all unihemispheric ROIs (BA44 ∩ BA45 ∩ Fo6&7 separately in each hemisphere) as well as the intersection of only the opercular and triangular parts of IFG (BA44 ∩ BA45 separately in each hemisphere) were computed. These intersection analyses aimed to identify the functional connectivity of the Broca’s complex in a broad sense including all opercular, triangular and orbital parts (Xiang et al., 2010) as well as that of the more conventionally recognized Broca’s area in a narrow sense including only the opercular and triangular parts (Amunts et al., 1999), in addition to that of their homologues in the right hemisphere. Finally, contrast/subtraction ALE analyses were performed to compare each ROI with the other two ROIs within the same hemisphere, following the logic of a previous MACM study of three subregions of the superior temporal sulcus (Erickson et al., 2017). Contrast analyses allowed better comparison of the ROIs since they exhibited overlap in coactivation and since the primary aim of the present study was to compare coactivation patterns of different IFG subdivisions. The following contrast analyses were performed: LBA44 > (LBA45 & LFo6&7), LBA45 > (LBA44 & LFo6&7), LFo6&7 > (LBA44 & LBA45); RBA44 > (RBA45 & RFo6&7), RBA45 > (RBA44 & RFo6&7), RFo6&7 > (RBA44 & RBA45). In other words, coactivations of each pair of ROIs in each hemisphere were subtracted from the third ROI within the same hemisphere iteratively using the Contrast Datasets utility in GingerALE, yielding specific coactivation patterns for each ROI. It should be mentioned that the ALE subtraction analysis utilizes permutation significance testing that controls for differences in the number of papers in each comparison set (Eickhoff et al., 2011; Erickson et al., 2017). Given that GingerALE performs contrast analyses based on already thresholded (in the present case, cluster-level FWE corrected) single-dataset images, and that the only thresholding available for contrast analyses in the current version of GingerALE at the time of analysis is false discovery rate (FDR), which is no longer recommended for spatially smooth data such as brain activations (Chumbley & Friston, 2009; Eickhoff et al., 2012), an uncorrected threshold of *p* < 0.05 with an extent threshold (minimum cluster size) of 100mm^3^ was applied for the contrast analyses.

For both the single ROI coactivation analyses and the contrast analyses, anatomical labels were generated as the nearest gray matter within 5mm for the activation peaks by the Talairach Daemon embedded in GingerALE (Lancaster et al., 1997; Lancaster et al., 2000). The results of the performed meta-analyses were visualized using the Mango software (Lancaster et al., 2010) and overlaid on the MNI template (Colin27_T1_seg_MNI.nii) available on GingerALE’s website. The foci extracted from each experiment using Sleuth and entered in the meta-analyses and the GingerALE outputs for each meta-analysis are accessible at https://www.doi.org/10.17632/nyx2gz9yww.1.

### Behavioral analysis

A behavioral analysis was performed using the Mango Behavioral Analysis Plugin (Lancaster et al., 2012), which was used in previous research to investigate functional specialization in different brain regions (Erickson et al., 2017; Sundermann & Pfleiderer, 2012). Utilizing the metadata of articles in the BrainMap database, the plugin calculates the observed fraction of activation coordinates for a given behavioral subdomain (e.g., cognition.attention, or cognition.language.phonology) that fall within a prespecified ROI and compares it to the fraction that would be expected if the distribution was random. If the difference between the observed and the expected fraction is high, then the ROI is associated with this behavior. The statistical threshold for significance was determined as a Z-score ≥ 3.0, which corresponds to a one-tailed (testing only positive association) *p*-value of .05 Bonferroni corrected for 51 behavioral subdomains in BrainMap (Lancaster et al., 2012). However, since the purpose of the behavioral analysis conducted in the present study was to examine specific linguistic functions of each ROI, rather than testing their domain-specificity (e.g., language vs. memory), the behavioral analysis was restricted to the phonology, semantics, speech and syntax subdomains, which are central to the neurocognitive models of language processing reviewed above. Hence, in the present study, the statistical threshold was set as Z > 2.24 (corresponding to a one-tailed *p*-value of .05 Bonferroni corrected for the four subdomains examined). The behavioral analysis was conducted on the same date as the database search using the same ROIs used for the MACM analysis as described above.

## Results

A chi-square test was conducted to examine whether the number of papers and experiments identified for each ROI significantly differed between the left and right hemispheres. It was found that significantly more papers and experiments were identified for each left-hemispheric ROI compared to the right-hemispheric ones (BA44: *X*^2^ > 26.60, *p* < .001; BA45: *X*^2^ > 24.82, *p* < .001; Fo6&7: *X*^2^ > 5.23, *p* < .022). This finding shows that more studies in the BrainMap database reported activations in the left IFG than the right IFG for language tasks.

### Coactivations of left-hemispheric ROIs

The coactivation and intersection results for each LIFG ROIs are visualized in Figure 2 and the coactivation results are summarized in Table 3. A more widespread coactivation pattern was observed for LBA44 (9 clusters, 29 peaks) than LBA45 (6 clusters, 17 peaks) and LFo6&7 (4 clusters, 11 peaks). Each ROI coactivated with itself most strongly. All LIFG ROIs significantly coactivated with ipsilateral regions within the frontal lobe and the temporal lobe (as shown in white in the lower panel of Figure 2), whereas only LBA44 showed additional coactivation with multiple parietal lobe structures. For all LIFG ROIs, the coactivations within the temporal lobe included BA22, which is commonly considered as Wernicke’s area, albeit with differences among the ROIs in terms of the cluster size. A large overlap of coactivation was observed between LBA44 and LBA45 especially in the frontal lobe ventrally and dorsally, but also in the left insular and temporal cortices. Although LIFG ROIs exhibited a strongly left-lateralized functional network, several right-hemispheric coactivations were also observed, especially for LBA44 (3 clusters and 10 peaks in the right cerebral hemisphere), to a lesser extent for LBA45 (2 clusters, 3 peaks) and none for LFo6&7 (no clusters). The right-hemispheric regions coactivating with both LBA44 and LBA45 include locations in the frontal lobe and the cingulate and insular cortices, while only LBA44 significantly coactivated with the right parietal cortex. Finally, several subcortical coactivations were identified. Only LBA44 showed coactivation in the thalamus, the basal ganglia (putamen) and the claustrum. Although all LIFG ROIs coactivated with parts of the cerebellum, the coactivation peaks for LBA45 and LFo6&7 were parts of a larger cluster together with predominantly fusiform gyrus involvement in the left temporal lobe, while LBA44 coactivated with the right cerebellum (culmen) specifically.

**Figure 2.**
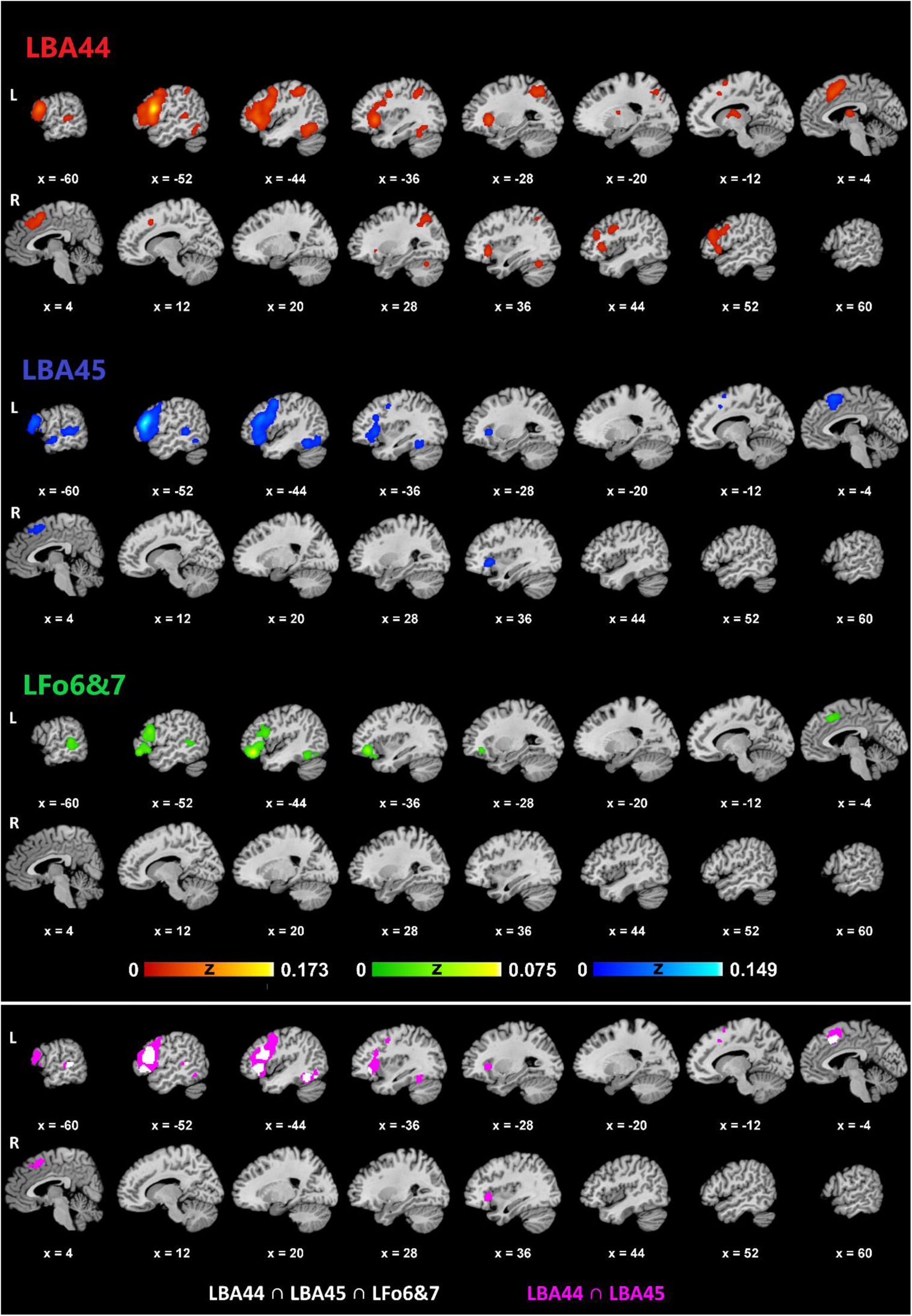
Coactivation results for the left-hemispheric ROIs [The upper panel shows the coactivation results for LBA44, LBA45 and LFo6&7 in the left and right hemispheres. The lower panel illustrates the intersection of LBA44, LBA45 and LFo6&7 (white) and of LBA44 and LBA45 (pink).

**Table 3.** Coactivation results for the left-hemispheric ROIs.

| Anatomical Label<br>(Nearest Gray Matter within 5mm) | BA | MNI Coordinates |  |  | ALE | Z | Cluster Size<br>(mm<sup>3</sup>) |
| --- | --- | --- | --- | --- | --- | --- | --- |
|  |  | x | y | z |  |  |  |
| LBA44 |  |  |  |  |  |  |  |
| L IFG | 44 | -54 | 12 | 18 | 0.173 | 16.60 | 41640 |
| L Claustrum |  | -32 | 22 | 0 | 0.087 | 9.96 |  |
| L IFG | 13 | -46 | 28 | 0 | 0.069 | 8.36 |  |
| L IFG | 46 | -46 | 32 | 10 | 0.061 | 7.60 |  |
| L Precentral Gyrus | 6 | -46 | 4 | 48 | 0.049 | 6.42 |  |
| L MFG | 10 | -42 | 46 | 18 | 0.025 | 3.62 |  |
| L FGmed | 32 | -4 | 16 | 46 | 0.058 | 7.34 | 11144 |
| L FGmed | 6 | -2 | 10 | 50 | 0.055 | 7.00 |  |
| R Cingulate Gyrus | 32 | 8 | 18 | 42 | 0.040 | 5.46 |  |
| R Cingulate Gyrus | 32 | 4 | 24 | 36 | 0.034 | 4.83 |  |
| R FGmed | 8 | 6 | 34 | 40 | 0.029 | 4.18 |  |
| L Cingulate Gyrus | 32 | -6 | 24 | 30 | 0.023 | 3.35 |  |
| R Insula |  | 36 | 24 | -8 | 0.046 | 6.18 | 10672 |
| R MFG | 46 | 50 | 28 | 16 | 0.044 | 5.86 |  |
| R IFG | 13 | 42 | 24 | 2 | 0.043 | 5.78 |  |
| R MFG | 9 | 50 | 30 | 24 | 0.041 | 5.63 |  |
| R IFG | 9 | 48 | 8 | 28 | 0.039 | 5.33 |  |
| R Insula | 13 | 50 | 16 | -4 | 0.037 | 5.19 |  |
| L IPL | 40 | -44 | -42 | 48 | 0.047 | 6.25 | 9248 |
| L Precuneus | 7 | -26 | -64 | 46 | 0.043 | 5.81 |  |
| L SPL | 7 | -28 | -56 | 46 | 0.043 | 5.79 |  |
| L IPL | 40 | -38 | -46 | 42 | 0.035 | 4.91 |  |
| L Fusiform Gyrus | 37 | -44 | -58 | -14 | 0.051 | 6.69 | 6224 |
| L Medial Dorsal Nucleus<br>(Thalamus) |  | -6 | -10 | 12 | 0.040 | 5.49 | 2400 |
| L Putamen (Lentiform Nucleus) |  | -12 | 2 | 6 | 0.023 | 3.34 |  |
| R SPL | 7 | 30 | -58 | 46 | 0.038 | 5.21 | 1960 |
| L MTG | 22 | -60 | -32 | 4 | 0.031 | 4.43 | 1816 |
| L STG | 22 | -54 | -40 | 8 | 0.030 | 4.34 |  |
| R Culmen (Cerebellum) |  | 38 | -60 | -26 | 0.033 | 4.70 | 1352 |
| LBA45 |  |  |  |  |  |  |  |
| L IFG | 45 | -52 | 28 | 14 | 0.149 | 15.29 | 36040 |
| L Precentral Gyrus | 6 | -44 | 2 | 46 | 0.057 | 7.49 |  |
| L IFG | 9 | -44 | 8 | 28 | 0.052 | 6.99 |  |
| L IFG | 47 | -44 | 26 | -14 | 0.049 | 6.63 |  |
| L Insula | 13 | -34 | 24 | 0 | 0.048 | 6.49 |  |
| L IFG | 47 | -46 | 22 | -10 | 0.047 | 6.39 |  |
| L Precentral Gyrus | 44 | -54 | 12 | 4 | 0.039 | 5.50 |  |
| L SFG | 6 | -4 | 16 | 56 | 0.049 | 6.60 | 6792 |
| L FGmed | 32 | -6 | 18 | 44 | 0.037 | 5.32 |  |
| R SFG | 8 | 6 | 30 | 48 | 0.024 | 3.62 |  |
| R Cingulate Gyrus | 32 | 6 | 26 | 40 | 0.021 | 3.25 |  |
| L Fusiform Gyrus | 37 | -44 | -52 | -18 | 0.046 | 6.27 | 5056 |
| L Declive (Cerebellum) |  | -46 | -70 | -14 | 0.032 | 4.74 |  |
| L MTG | 22 | -56 | -38 | 2 | 0.045 | 6.22 | 3608 |
| R Insula |  | 38 | 22 | -4 | 0.040 | 5.63 | 1840 |
| L MTG | 21 | -58 | -10 | -12 | 0.033 | 4.87 | 1400 |
| L MTG | 21 | -58 | -2 | -18 | 0.028 | 4.18 |  |

| LFo6&7 |  |  |  |  |  |  |  |
| --- | --- | --- | --- | --- | --- | --- | --- |
| L MFG | 47 | -46 | 36 | -14 | 0.075 | 10.75 | 15440 |
| L IFG | 9 | -50 | 18 | 22 | 0.037 | 6.64 |  |
| L IFG | 47 | -50 | 24 | -8 | 0.027 | 5.30 |  |
| L IFG | 45 | -50 | 26 | 14 | 0.026 | 5.14 |  |
| L IFG | 47 | -38 | 24 | -20 | 0.015 | 3.47 |  |
| L MTG | 22 | -62 | -38 | 4 | 0.027 | 5.21 | 3168 |
| L MTG | 21 | -60 | -38 | -2 | 0.023 | 4.72 |  |
| L MTG | 22 | -54 | -48 | 2 | 0.019 | 4.04 |  |
| L Fusiform Gyrus | 37 | -46 | -52 | -18 | 0.024 | 4.77 | 1480 |
| L Declive (Cerebellum) |  | -46 | -68 | -18 | 0.016 | 3.61 |  |
| L FGmed | 6 | -6 | 20 | 44 | 0.024 | 4.81 | 1400 |

Note: MNI Coordinates correspond to cluster peaks. L: Left, R: Right, FGmed: Medial frontal gyrus, IFG: Inferior frontal gyrus, IPL: Inferior parietal lobule, ITG: Inferior temporal gyrus, MFG: Middle frontal gyrus, MTG: Middle temporal gyrus, SFG: Superior frontal gyrus, SPL: Superior parietal lobule, STG: Superior temporal gyrus.

### Contrasts among left-hemispheric ROIs

As illustrated in Figure 3 and detailed in Table 4, the contrast/subtraction results revealed ROI-specific coactivations when the coactivation network of each ROI pair was subtracted from the other ROI within the same hemisphere. Similar to the main coactivation results summarized above, each LIFG ROI coactivated with itself, thereby revealing a coactivation pattern in the left IFG reminiscent of the predefined ROI boundary. Again, a more extensive coactivation contrast network was identified for LBA44 (14 clusters, 29 peaks) than LBA45 (3 clusters, 3 peaks) and LFo6&7 (3 clusters, 8 peaks). In the left hemisphere, LBA44 exhibited a more robust dorsal coactivation pattern extending into the precentral gyrus in the frontal lobe and into the inferior and superior parietal lobule including the precuneus. LBA45 and LFo6&7, on the other hand, revealed a more ventral distribution spanning only the frontal and temporal lobes. Within the temporal lobe, LBA44 showed specific coactivation solely in the fusiform gyrus, except for a single peak at STG which was a tiny part (1.6%) of a much larger frontal cluster. However, LBA45 and LFo6&7 coactivated more specifically with the anterior and posterior MTG, respectively. Also, only LBA44 revealed specific coactivation in the left insula, while only LFo6&7 strongly coactivated with the left cingulate gyrus. As for the coactivation patterns in the right hemisphere, only LBA44 showed specific right-hemispheric coactivations in the frontal lobe (middle and superior frontal gyri, precentral gyrus), parietal lobe (precuneus) and the cingulate gyrus. The extensive coactivation network specific to LBA44 also spanned left-hemispheric subcortical structures (claustrum, thalamus) and the culmen in the right cerebellar hemisphere.

**Figure 3.**
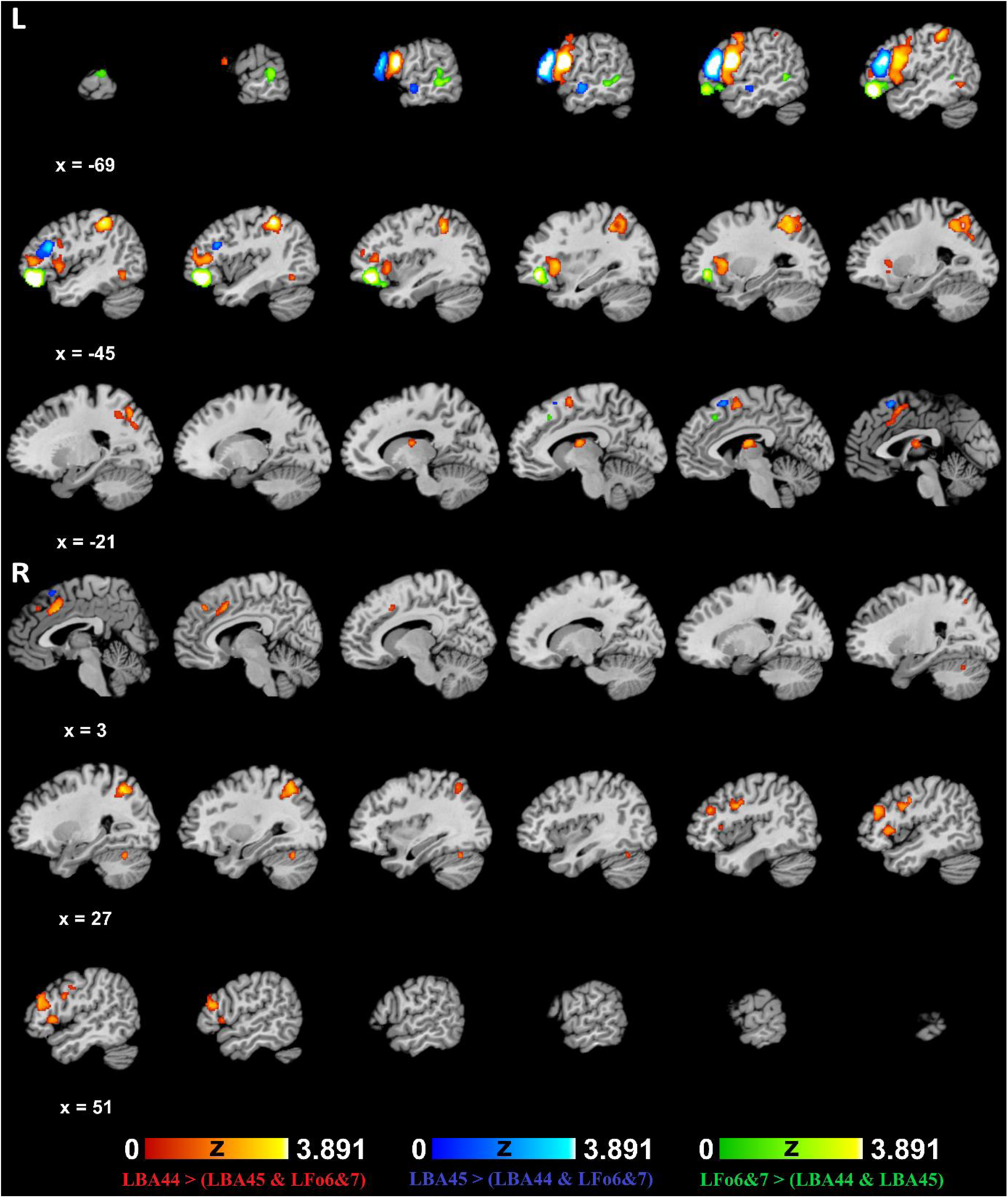
Contrast results for the left-hemispheric ROIs.

**Table 4.**
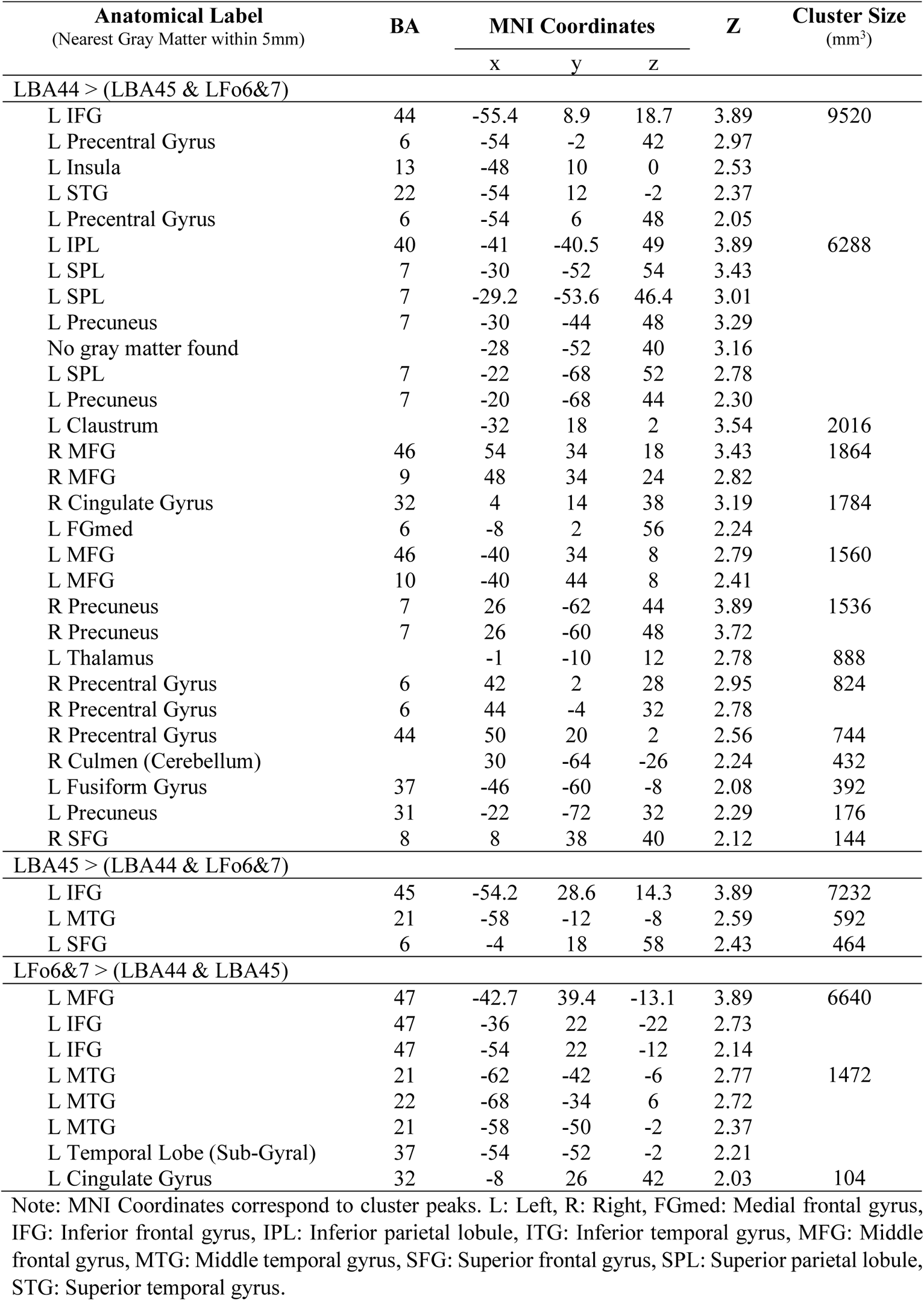
Contrast results for the left-hemispheric ROIs.

### Coactivations of right-hemispheric ROIs

Coactivation results for RIFG ROIs are shown in Figure 4 and Table 5. Similar to the LIFG results, the right-hemispheric ROIs strongly coactivated with themselves and adjacent regions. Also in parallel with the LIFG results, the opercular portion of RIFG; i.e., RBA44, revealed a more widespread coactivation pattern (9 clusters, 16 peaks) than RBA45 (6 clusters, 13 peaks) and RFo6&7 (3 clusters, 6 peaks). Unlike LIFG results, however, there was almost no overlap among all three ROIs in either the left or the right hemisphere, while RBA44 and RBA45 exhibited coactivation overlap in the left and right frontal and insular cortices. Within the right hemisphere, RIFG ROIs coactivated almost exclusively with frontal regions, except for the right insular coactivation for RBA44 and RBA45, and for the coactivation in the right anterior STG, which was part of a larger frontal cluster, for RBA44. In the contralateral left hemisphere, on the other hand, more widespread coactivation was observed for all RIFG ROIs. RBA44 and RBA45 also coactivated with the parietal lobe (different sites within the inferior parietal lobule). All RIFG ROIs involved homotopic functional connectivity with the left-hemispheric mirror regions, which extended into surrounding regions within IFG and MFG (for all RIFG ROIs), the precentral gyrus and the medial frontal gyrus (for RBA44 and RBA45) and SFG (for RBA44). All right-hemispheric ROIs also coactivated with the posterior temporal regions, with RBA44 exhibiting functional connectivity with a more superior site (MTG) than RBA45 and RFo6&7 (fusiform gyrus). In addition, RBA44 was associated with significant coactivation in the left insular and cingulate cortices. Finally, subcortical coactivations were identified for RBA44 (left thalamus) and RBA45 (left and right claustra).

**Figure 4.**
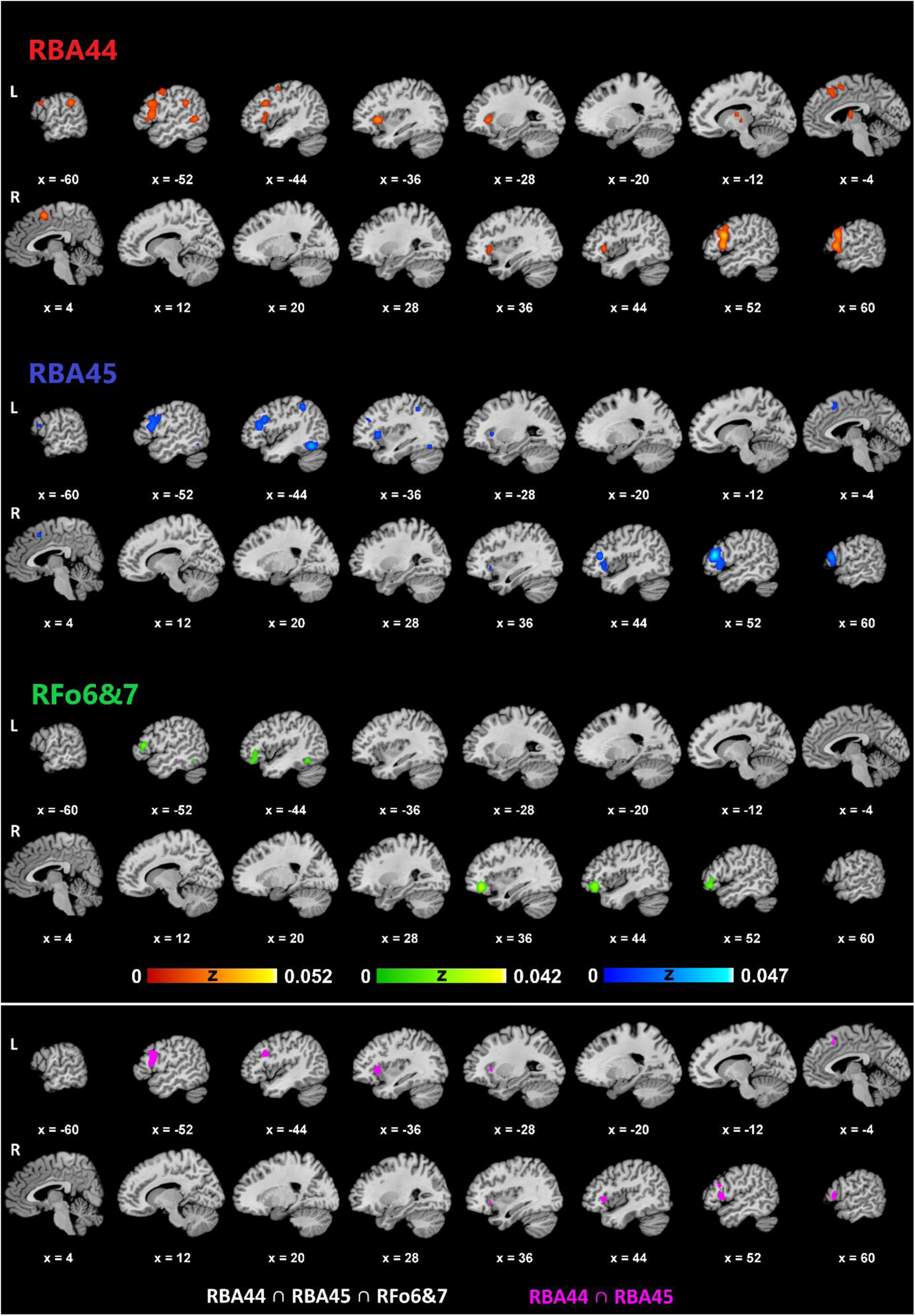
Coactivation results for the right-hemispheric ROIs. [The upper panel shows the coactivation results for RBA44, RBA45 and RFo6&7 in the left and right hemispheres. The lower panel illustrates the intersection of RBA44, RBA45 and RFo6&7 (white), which revealed only a tiny blob in the left BA45 (at around x = −52, y = 23, z = 11) and the intersection of RBA44 and RBA45 (pink).

**Table 5.** Coactivation results for the right-hemispheric ROIs.

| Anatomical Label<br>(Nearest Gray Matter within 5mm) | BA | MNI Coordinates |  |  | ALE | Z | Cluster Size<br>(mm<sup>3</sup>) |
| --- | --- | --- | --- | --- | --- | --- | --- |
|  |  | x | y | z |  |  |  |
| RBA44 |  |  |  |  |  |  |  |
| L Insula | 13 | -36 | 20 | 0 | 0.030 | 6.01 | 8000 |
| L IFG | 44 | -50 | 16 | 10 | 0.025 | 5.31 |  |
| L IFG | 9 | -50 | 16 | 28 | 0.023 | 4.97 |  |
| R IFG | 9 | 54 | 14 | 22 | 0.052 | 8.81 | 7400 |
| R Precentral Gyrus | 44 | 56 | 16 | 6 | 0.040 | 7.29 |  |
| R MFG | 9 | 56 | 10 | 34 | 0.022 | 4.81 |  |
| R STG | 22 | 54 | 8 | -10 | 0.014 | 3.48 |  |
| L IPL | 40 | -56 | -40 | 30 | 0.030 | 5.97 | 1488 |
| L FGmed | 6 | 2 | 6 | 54 | 0.026 | 5.42 | 1472 |
| L Precentral Gyrus | 4 | -48 | -4 | 52 | 0.021 | 4.64 | 1088 |
| R Insula | 13 | 42 | 22 | 0 | 0.021 | 4.60 | 1008 |
| L Cingulate Gyrus | 32 | -2 | 22 | 42 | 0.017 | 4.00 | 1000 |
| L SFG | 8 | -4 | 24 | 48 | 0.017 | 3.99 |  |
| L Anterior Nucleus (Thalamus) |  | -8 | -8 | 10 | 0.021 | 4.72 | 968 |
| L Mammillary Body (Thalamus) |  | -10 | -16 | 0 | 0.014 | 3.49 |  |
| L MTG | 37 | -52 | -56 | 2 | 0.021 | 4.69 | 784 |
| RBA45 |  |  |  |  |  |  |  |
| R MFG | 46 | 50 | 30 | 14 | 0.047 | 8.40 | 8224 |
| R IFG | 47 | 50 | 20 | -6 | 0.020 | 4.54 |  |
| R Insula | 13 | 42 | 22 | 2 | 0.016 | 4.00 |  |
| R Claustrum |  | 34 | 22 | -6 | 0.014 | 3.58 |  |
| L MFG | 46 | -46 | 28 | 14 | 0.022 | 4.84 | 5480 |
| L IFG | 44 | -52 | 16 | 14 | 0.021 | 4.77 |  |
| L MFG | 9 | -44 | 18 | 28 | 0.020 | 4.51 |  |
| L Precentral Gyrus | 6 | -52 | 4 | 32 | 0.014 | 3.50 |  |
| L IFG | 46 | -50 | 38 | 8 | 0.013 | 3.33 |  |
| L Fusiform Gyrus | 37 | -44 | -58 | -16 | 0.034 | 6.65 | 2712 |
| L Claustrum |  | -32 | 20 | 2 | 0.023 | 5.14 | 1152 |
| L IPL | 40 | -42 | -44 | 48 | 0.021 | 4.83 | 936 |
| L FGmed | 32 | 0 | 16 | 48 | 0.016 | 3.98 | 808 |
| RFo6&7 |  |  |  |  |  |  |  |
| R IFG | 47 | 40 | 36 | -12 | 0.042 | 8.00 | 4472 |
| R IFG | 46 | 50 | 34 | 6 | 0.011 | 3.22 |  |
| L IFG | 45 | -52 | 28 | 10 | 0.019 | 4.72 | 1816 |
| L MFG | 11 | -44 | 38 | -18 | 0.014 | 3.82 |  |
| L IFG | 47 | -44 | 32 | -12 | 0.013 | 3.72 |  |
| L Fusiform Gyrus | 37 | -48 | -54 | -16 | 0.021 | 5.06 | 792 |

Note: MNI Coordinates correspond to cluster peaks. L: Left, R: Right, FGmed: Medial frontal gyrus, IFG: Inferior frontal gyrus, IPL: Inferior parietal lobule, ITG: Inferior temporal gyrus, MFG: Middle frontal gyrus, MTG: Middle temporal gyrus, SFG: Superior frontal gyrus, SPL: Superior parietal lobule, STG: Superior temporal gyrus.

### Contrasts among right-hemispheric ROIs

The contrast results for the right-hemispheric ROIs are shown in Figure 5 and Table 6. In line with the main coactivation results, each ROI exhibited specific coactivation with itself. As before, a more extensive coactivation contrast network was identified for RBA44 (8 clusters, 17 peaks) than RFo6&7 (5 clusters, 9 peaks) and RBA45 (4 clusters, 7 peaks). In the right-hemisphere, region-specific coactivation was restricted to the frontal lobe, with each right-hemispheric ROI specifically coactivating with itself, and with a few other frontal sites including the precentral gyrus (for RBA44), medial frontal gyrus (for RBA44) and SFG (for RFo6&7). In the left hemisphere, all right-hemispheric ROIs showed specific coactivation within the frontal and temporal lobes, while only RBA44 and RBA45 specifically coactivated with the parietal lobe (inferior parietal lobule), which was much stronger for RBA44. In the left frontal lobe, the right-hemispheric ROIs exhibited specific homotopic coactivation in and around LIFG (for RBA44 and RFo6&7), MFG (for RBA45 and RFo6&7), the precentral gyrus (for RBA44) and the medial frontal gyrus (for RFo6&7). In the left temporal lobe, RBA45 and RFo6&7 specifically coactivated with the fusiform gyrus, whereas RBA44 showed specific coactivation more superiorly within MTG. In addition, only RBA44 exhibited specific coactivation in the left insula. The right-hemispheric ROIs also revealed specific subcortical coactivation in the left claustrum (for RBA44) and the left cerebellar hemisphere (declive for RBA45 and culmen for RFo6&7), the latter being parts of larger clusters in the left fusiform gyrus.

**Figure 5.**
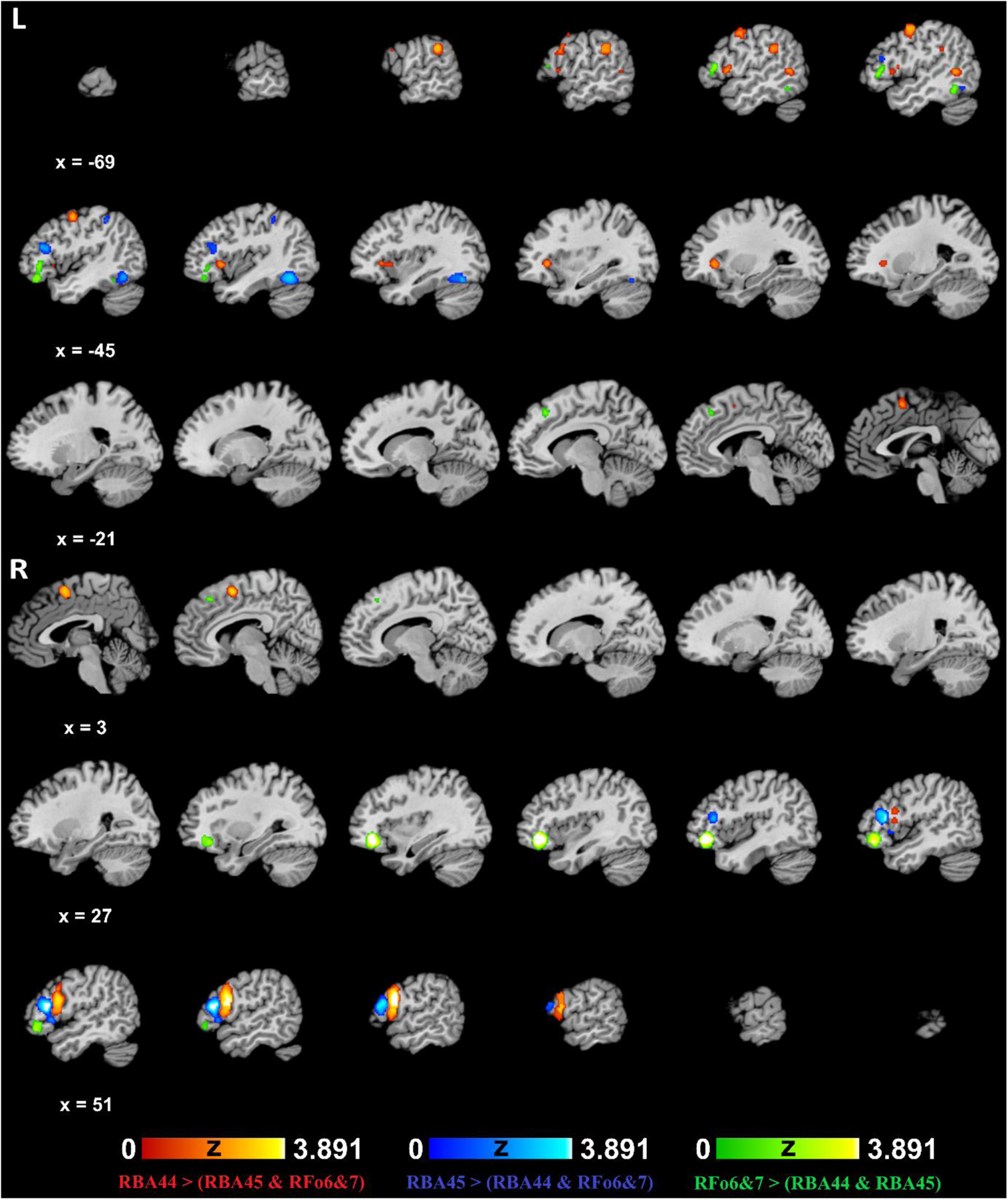
Contrast results for the right-hemispheric ROIs.

**Table 6.** Contrast results for the right-hemispheric ROIs.

| Anatomical Label<br>(Nearest Gray Matter within 5mm) | BA | MNI Coordinates |  |  | Z | Cluster Size<br>(mm<sup>3</sup>) |
| --- | --- | --- | --- | --- | --- | --- |
|  |  | x | y | z |  |  |
| RBA44 > (RBA45 & RFo6&7) |  |  |  |  |  |  |
| R IFG | 44 | 58.2 | 10.6 | 20 | 3.89 | 4856 |
| R Precentral Gyrus | 44 | 59 | 12 | 8 | 3.54 |  |
| L IPL | 40 | -62 | -38 | 30 | 2.86 | 1056 |
| L IPL | 40 | -62 | -34 | 29 | 2.82 |  |
| L IPL | 40 | -52 | -36 | 32 | 2.14 |  |
| R FGmed | 6 | 7 | 4 | 56 | 3.12 | 1016 |
| L Precentral Gyrus | 4 | -50 | -2 | 54 | 2.69 | 984 |
| L Precentral Gyrus | 4 | -50 | -6 | 54 | 2.64 |  |
| L Precentral Gyrus | 4 | -52 | -4 | 42 | 2.34 |  |
| L Insula |  | -42 | 16 | -2 | 2.26 | 880 |
| L IFG | 45 | -36 | 28 | 0 | 2.18 |  |
| L Claustrum |  | -26 | 26 | 0 | 2.12 |  |
| L ITG | 37 | -48 | -56 | 2 | 2.31 | 408 |
| L MTG | 37 | -52 | -54 | 6 | 2.30 |  |
| L Precentral Gyrus | 44 | -52 | 10 | 8 | 2.08 | 296 |
| L IFG | 9 | -58 | 14 | 26 | 1.91 | 208 |
| L Precentral Gyrus | 6 | -58 | 10 | 34 | 1.90 |  |
| RBA45 > (RBA44 & RFo6&7) |  |  |  |  |  |  |
| R IFG | 45 | 52.6 | 25.6 | 13 | 3.89 | 3960 |
| R IFG | 45 | 52 | 20 | -2 | 2.29 |  |
| L Declive (Cerebellum) |  | -38 | -61 | -16 | 3.35 | 1504 |
| L Fusiform Gyrus | 37 | -38 | -55 | -14 | 3.24 |  |
| L MFG | 46 | -44 | 26 | 18 | 2.58 | 584 |
| L MFG | 46 | -46 | 30 | 18 | 2.52 |  |
| L IPL | 40 | -46 | -40 | 48 | 2.04 | 128 |
| RFo6&7 > (RBA44 & RBA45) |  |  |  |  |  |  |
| R IFG | 47 | 39.6 | 36.9 | -11.7 | 3.89 | 3936 |
| L MFG | 47 | -42 | 38 | -14 | 2.30 | 1072 |
| L IFG | 45 | -54 | 28 | 8 | 2.26 |  |
| L IFG | 13 | -48 | 32 | 0 | 2.13 |  |
| L MFG | 47 | -44 | 36 | -6 | 2.04 |  |
| L Culmen (Cerebellum) |  | -50 | -52 | -20 | 2.06 | 248 |
| L Fusiform Gyrus | 37 | -52 | -54 | -16 | 2.05 |  |
| L FGmed | 8 | -8 | 32 | 44 | 2.37 | 200 |
| R SFG | 8 | 8 | 30 | 48 | 2.64 | 104 |

Note: MNI Coordinates correspond to cluster peaks. L: Left, R: Right, FGmed: Medial frontal gyrus, IFG: Inferior frontal gyrus, IPL: Inferior parietal lobule, ITG: Inferior temporal gyrus, MFG: Middle frontal gyrus, MTG: Middle temporal gyrus, SFG: Superior frontal gyrus, SPL: Superior parietal lobule, STG: Superior temporal gyrus.

### Behavioral Analysis Results

As shown in Figure 6, LBA44 was significantly associated with phonology, semantics and speech, while its association with syntax (Z = 2.15) was slightly below the statistical threshold (Z > 2.24). Also, LBA45 was only significantly related with semantics. None of the other ROIs had significant associations with the behavioral subdomains examined.

**Figure 6.**
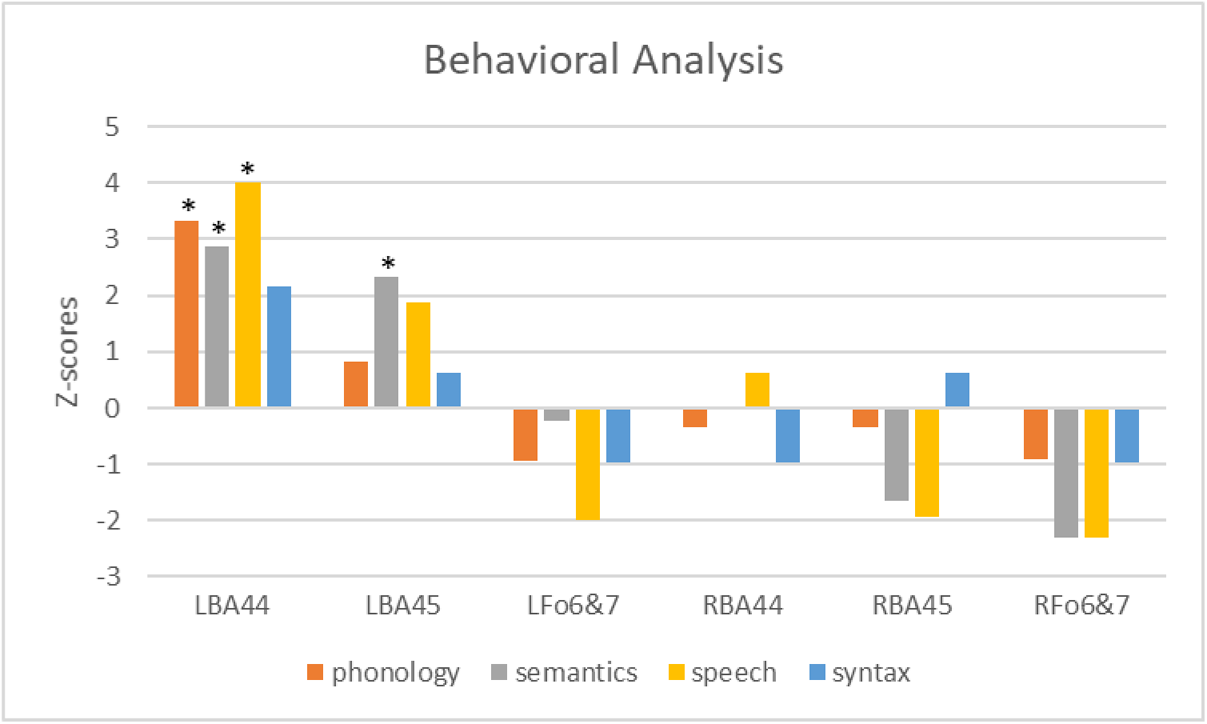
Behavioral analysis results for each ROI [* Bonferroni-corrected *p* < 0.05].

## Discussion

The current study investigated language-related coactivation patterns of three cytoarchitectonically-defined subdivisions (pars opercularis / BA44, pars triangularis / BA45, and pars orbitalis / BA47 / Fo6&7) of the inferior frontal gyrus in the left and right hemispheres. A series of meta-analytic connectivity modeling analyses identified functional connectivity of these ROIs during language tasks and compared IFG subdivisions through subtraction analyses to reveal coactivation patterns specific to each ROI. The database search identified more studies reporting coactivations in the left-hemispheric ROIs compared to the right-hemispheric ones. Also, a more widespread and robust coactivation pattern was observed in the left hemisphere than the right for both left- and right-hemispheric ROIs, confirming left-hemispheric dominance of IFG functional connectivity for language. Another finding was that among both left and right hemispheric ROIs, BA44 revealed a more widespread coactivation pattern compared to BA45 and Fo6&7, highlighting this region as a core component of the functional network for language. The findings from the left- and right-hemispheric ROIs and their implications for neurocognitive models of language processing are discussed below.

### Coactivation patterns of left-hemispheric ROIs

The left-hemispheric ROIs coactivated more strongly and extensively with regions in the left hemisphere than with the right hemisphere. LBA44 revealed a more widespread, dorsal coactivation pattern spanning the left frontal, temporal and parietal cortices while the coactivation network of LBA45 and LFo6&7 was mostly restricted ventrally to the frontal and temporal regions. Similar findings were reported in a resting-state functional connectivity study comparing similarly defined IFG subdivisions (Xiang et al., 2010), which attributed this pattern of connectivity to inherent distinctions among the IFG subdivisions and their functional connections within the perisylvian language system. The involvement of LBA44 with a dorsal coactivation pattern and LBA45 and LFo6&7 with a ventral pattern is consistent with several neurocognitive models of language processing that emphasize ventral-dorsal disparity in the functional connectivity of Broca’s area (Friederici, 2002, 2011, 2012; Hagoort, 2013, 2016; Hickok & Poeppel, 2004, 2007; Poeppel et al., 2012; Xiang et al., 2010). In general, these models implicate posterior-dorsal LIFG usually comprising LBA44, but sometimes also LBA45, with dorsal connections within the left frontal, temporal and sometimes parietal cortices, and the anterior-ventral LIFG usually comprising LBA47, but sometimes also LBA45, with ventral connections within the left frontal and temporal cortices. The functions attributed to the dorsal network range from syntactic processing (Friederici, 2002, 2011, 2012), syntactic and phonological processing (Hagoort, 2013, 2016; Xiang et al., 2010) to speech production (Hickok & Poeppel, 2004, 2007; Poeppel et al., 2012). On the other hand, the ventral network has been associated with semantic processing (Friederici, 2002, 2011, 2012; Hagoort, 2013, 2016; Xiang et al., 2010) and speech comprehension (Hickok & Poeppel, 2004, 2007; Poeppel et al., 2012). These associations are partially corroborated by the behavioral analysis results, which revealed greatest and significant functional specificity in LBA44 for speech, phonology and semantics, whereas LBA45 exhibited significant functional specificity only for semantics. Furthermore, LBA44 was involved with syntax more than the other ROIs, although this involvement did not reach statistical significance. However, no functional specialization for language subdomains was found within LFo6&7, which was previously associated with semantic processing (Hagoort, 2013, 2016; Xiang et al., 2010). Overall, the behavioral analysis results support the neurocognitive models arguing for a functional segregation within LIFG for different linguistic functions (Friederici, 2002, 2011, 2012; Hagoort, 2013, 2016; Xiang et al., 2010).

In the left frontal lobe, LBA44 showed the greatest and most widespread coactivation, followed by LBA45 and finally by LFo6&7. Particularly, LBA44 and LBA45 coactivated with more extensive frontal regions spanning the inferior, middle and medial frontal gyri and the precentral gyrus as well as the insular cortex, whereas the coactivation of LFo6&7 was restricted largely to anterior-ventral frontal regions (middle and inferior frontal gyri), but also included the medial frontal gyrus. These findings are compatible with the Dual-Stream Model, which highlights the functional network involving dorsal LIFG (LBA44 and LBA45), the left premotor cortex and the left insula with articulation (Hickok & Poeppel, 2007; Poeppel et al., 2012). The coactivation of LBA44 with the left premotor cortex (BA6) is also consistent with the DP Model, which associates these regions with processing automatized procedural memories including rule-based grammatical processing (Ullman, 2016). In addition, the functional connectivity of LIFG with the left dorsolateral prefrontal cortex and the dorsomedial prefrontal cortex including the anterior cingulate cortex is compatible with the MUC Model, which implicates these regions with executive control processes including attention and cognitive control underlying language (Hagoort, 2005, 2013, 2016; Xiang et al., 2010). Notably, as shown by the contrast analyses, LBA44 specifically coactivated with the superior-posterior part of the dorsolateral prefrontal cortex, while LBA45 specifically coactivated with the more anterior part. The coactivation results also related LFo6&7 with the inferior dorsolateral prefrontal cortex. Taken together, this ventral-anterior-posterior gradient within the dorsolateral prefrontal cortex (LBA46, 9 and 6) and its corresponding functional connectivity with ventral-anterior-posterior LIFG may represent the control processes underlying semantic, syntactic and phonological processes in parallel with the claims of the MUC Model (Hagoort, 2005, 2013, 2016; Xiang et al., 2010).

Of the left-hemispheric ROIs, only LBA44 coactivated with regions within the parietal lobe, specifically in the inferior parietal lobule (IPL) including the supramarginal gyrus and the superior parietal lobule (SPL) including the precuneus. The coactivation in SPL was bilateral, whereas that of IPL was found only in the left hemisphere. SPL is a functionally heterogenous region that has been bilaterally involved in various multisensory processes including action and visuomotor processing, visual perception, spatial cognition, reasoning, working memory, and attention (Wang et al., 2015). IPL, on the other hand, has been more commonly implicated in language processing. Also known as Geschwind’s territory (Catani, Jones, & Ffytche, 2005; Geschwind, 2006), left IPL was shown in DTI studies to indirectly connect Broca’s and Wernicke’s areas in parallel with the arcuate fasciculus which provides a direct pathway between the two (Catani et al., 2005). Various cognitive functions have been attributed to IPL including semantic processing (Catani et al., 2005), syntactic processing (Hagoort & Indefrey, 2014), reading and language learning (Barbeau et al., 2017), phonological storage as part of working memory (Baldo & Cronkers, 2006; Vigneau et al., 2006), number processing (Arsalidou & Taylor, 2011; Hung et al., 2015), processing tools and object-directed actions (Chen, Garcea, Jacobs, & Mahon, 2018), among others, stressing its role in various functional domains as a major network hub (Igelström & Graziano, 2017). According to the Dual-Stream

Model, the temporoparietal region at the posterior end of the sylvian fissure (area Spt) is part of the dorsal articulatory network together with dorsal LIFG (BA44 and 45), the left premotor cortex and the left insula, and is responsible for sensorimotor interface; i.e., translation of linguistic information between sensory (auditory, visual etc.) and production (motor) systems (Hickok & Poeppel, 2007; Poeppel et al., 2012). The coactivation of LBA44 with IPL observed here may be interpreted along these lines, given also that significant functional specificity for speech was found in LBA44 in the behavioral analysis. However, the IPL coactivation with LBA44 was observed more superiorly than area Spt; hence, it is not clear to what extent this coactivation converges with the functions attributed to area Spt. Functional connectivity of LBA44 and left IPL is also consistent with a recent version of Friederici’s Model, which argues for a second dorsal pathway connecting posterior STG with the premotor cortex, in addition to the primary dorsal pathway between LBA44 and posterior STG (Friederici, 2011, 2012). This second dorsal pathway is further divided into direct and indirect routes, the direct route connecting posterior STG directly with the premotor cortex, and the indirect route connecting posterior STG with the premotor cortex via the inferior parietal cortex. The premotor cortex in turn makes a loop with LBA44. It is argued that this second dorsal pathway facilitates mapping between phonological and motor/articulatory information. The coactivation of LBA44 with left IPL (supramarginal gyrus) is also consistent with the predictions of the MUC Model, which associates this connectivity with phonological processing (Hagoort, 2013, 2016; Xiang et al., 2010), given that left IPL is involved in phonological short-term memory. However, this model also predicts left parietal coactivations for LBA45 and LFo6&7 for syntactic and semantic processing, respectively, which were absent in the present study. Overall, these three neurocognitive models of language processing converge on the importance of the connection between the left inferior parietal cortex and dorsal LIFG for speech and phonology, which is also consistent with the functional specificity for speech and phonology observed in LBA44 in the behavioral analysis.

Within left temporal lobe, the coactivation networks of the three left-hemispheric ROIs converged in the posterior temporal lobe; specifically, on the posterior portion of the left middle temporal gyrus overlapping with portions of BA22; i.e., Wernicke’s area. The contrast analyses, however, showed that only LFo6&7 specifically coactivated with parts of this posterior temporal region. The posterior superior and middle temporal lobe has been associated with various linguistic functions, primarily including phonological processing (particularly STG) (Turkeltaub & Branch Coslett, 2010; Vigneau et al., 2006), syntactic processing (Hagoort & Indefrey, 2014; Heard & Lee, 2020; Rodd et al., 2015; Vigneau et al., 2006; Walenski et al., 2019), semantic processing (Hagoort & Indefrey, 2014; Rodd et al., 2015; Vigneau et al., 2006), and sentence comprehension (Vigneau et al., 2006; Walenski et al., 2019). This multifunctional nature of the left posterior temporal lobe may have contributed to the present observation of convergent coactivation for all LIFG subdivisions in that area.

Another region where significant functional connectivity was found is the anterior middle temporal gyrus, which significantly coactivated with only LBA45. The anterior temporal lobe has been associated with sentence comprehension in an ALE meta-analysis (Walenski et al., 2019), and several neuroimaging studies implicated this region in combinatorial processing at the sentence level (higher syntactic and compositional semantic processing), usually identified through sentence versus word list contrasts, (Brennan et al., 2012; Brennan & Pylkkänen, 2012; Bulut, Hung, Tzeng, & Wu, 2017; Humphries, Willard, Buchsbaum, & Hickok, 2001; Colin Humphries, Love, Swinney, & Hickok, 2005; Rogalsky & Hickok, 2009). In light of these previous studies and of the behavioral analysis which revealed significant functional specificity in LBA45 only for semantics, it could be argued that functional connectivity between LBA45 and anterior MTG underlies semantic processing, possibly at the sentence level. This finding is consistent with the Dual-Stream Model, which associates the left anterior temporal lobe with combinatorial processing of syntactic and semantic information, and conceptualizes a direct connection between the ventral and dorsal streams via the left anterior temporal lobe and dorsal LIFG (LBA45, but also LBA44) (Hickok & Poeppel, 2004, 2007; Poeppel et al., 2012). LBA45 – left anterior temporal lobe coactivation is also compatible with Friederici’s Modal, according to which anterior STG, MTG and anterior IFG (BA45, but also BA47) constitute the ventral pathway underlying semantic processing (Friederici, 2002, 2011, 2012).

Another area that exhibited convergent coactivation for all left-hemispheric ROIs within the temporal lobe was the left fusiform gyrus. However, the contrast analyses showed that only LBA44 specifically coactivated with parts of this region. The left fusiform gyrus has been associated with reading and recognition of visual word forms (McCandliss, Cohen, & Dehaene, 2003), as well as with recognition of meaningful visual objects in general (Devlin, Jamison, Gonnerman, & Matthews, 2006). In particular, neuroimaging research as well as clinical studies of semantic dementia involved this region with lexical-semantic processing (Ardila, Bernal, & Rosselli, 2015; Binder, Desai, Graves, & Conant, 2009; Ding et al., 2016; Wheatley, Weisberg, Beauchamp, & Martin, 2005). Given that LBA44 and LBA45 exhibited stronger coactivation with this region than LFo6&7, and that only the former two ROIs revealed functional specialization for semantic processing in the behavioral analysis, it is likely that LIFG – left fusiform gyrus coactivation may underlie semantic processing, possibly at the lexical level. The neurocognitive models of language processing do not explicitly associate the fusiform gyrus in the language network, except for involvement by the MUC Model of the LBA47 – left posterior inferior temporal gyrus connection with semantic processing (Hagoort, 2005, 2013, 2016). In the present study, though, the coactivation was within the fusiform gyrus, which is on the basal surface of the temporal lobe. Also, not only LFo6&7 corresponding to LBA47, but all three LIFG subdivisions significantly coactivated with the fusiform gyrus, contrary to the model’s prediction of mainly LBA47 involvement with the left inferior temporal lobe.

Although LIFG ROIs revealed a strongly left-lateralized functional network, several right-hemispheric coactivations were also observed, especially for LBA44, to a lesser extent for LBA45, but none for LFo6&7. The right-hemispheric regions coactivating with both LBA44 and LBA45 include locations in the frontal lobe and the cingulate and insular cortices, while only LBA44 significantly coactivated with the right parietal cortex. Moreover, contrast analyses revealed that only LBA44 showed specific coactivation with right-hemispheric structures in the frontal lobe (MFG, SFG, precentral gyrus also spanning some homotopic RIFG territory), parietal lobe (precuneus) and the cingulate gyrus. While the left hemisphere has generally been considered as the dominant hemisphere in language processing, the linguistic role of the right hemisphere has been controversial. Some of the communicative functions attributed in neuroimaging research to the right hemisphere include figurative and metaphorical language processing (Gainotti, 2016, but cf. Cardillo, McQuire, & Chatterjee, 2018), processing contextual and coherent meaning particularly in discourse (Vigneau et al., 2011; Xu, Kemeny, Park, Frattali, & Braun, 2005; Zempleni et al., 1998) and emotional/affective prosody (George et al., 1996; Patel et al., 2018). In parallel with these associations, lesions in the right hemisphere have been reported to involve deficits in pragmatics; i.e., context-dependent use of language and nonverbal elements in communication (Lundgren & Brownell, 2016; Lundgren, Brownell, Cayer-Meade, Milione, & Kearns, 2011; Parola et al., 2016), and emotional prosody (Dara, Bang, Gottesman, & Hillis, 2014; Ross & Mesulam, 1979; Ross & Monnot, 2008; Stockbridge et al., 2021).

Although it is difficult to characterize specific contributions of the right-hemispheric coactivations observed in the present study, they are unlikely to be merely due to contextual, prosodic and pragmatic processes, since LBA44, which showed greatest right-hemispheric coactivation, exhibited functional specificity for phonology, semantics and speech, and, to a limited extent, for syntax, and since the experiments included in the MACM analyses of LBA44 involved manipulations at various linguistic levels; i.e., speech, semantics, phonology, syntax and orthography. The role of the right hemisphere in the language network is discussed further in the following section. The language network proposed by the models tested here is strongly left-lateralized, with only limited involvement of the right hemisphere. Although the Dual-Stream Model recognizes some right-hemispheric contribution to language processing, it claims that only the ventral stream underlying speech comprehension involves bilateral temporal cortices, whereas the dorsal stream responsible for speech production is left dominant (Hickok & Poeppel, 2004, 2007; Poeppel et al., 2012). Therefore, the present finding of especially LBA44 coactivation within the right hemisphere challenges all the neurocognitive models of language processing addressed here.

Previous research associated subcortical structures primarily including the thalamus and the basal ganglia with language; however, how these structures contribute to language processing and how they are connected to the language network remains to be elucidated (Friederici, 2012). In the present study, subcortical coactivations were seen exclusively for LBA44 in the thalamus, the basal ganglia (putamen) and the claustrum in the left hemisphere as part of a single cluster in the main coactivation analysis. This cluster predominantly included the thalamus (96.7%), while overlapping with fragments of the caudate body (1.9%) and the putamen (1.4%). Neuroimaging studies, animal models and neurodegenerative disorders of the basal ganglia including the caudate nucleus and the putamen associated this structure with various cognitive, motor and emotional processes particularly including reward processing, learning and memory, and motor control (Albin, Young, & Penney, 1989; Chakravarthy, Joseph, & Bapi, 2010; Groenewegen, 2003; Lanciego, Luquin, & Obeso, 2012; Packard & Knowlton, 2002). Indeed, the basal ganglia-thalamo-cortical system is associated with a wide spectrum of sensorimotor, cognitive, emotional and motivational brain functions (Alexander & Crutcher, 1990; Groenewegen, 2003). An MACM study of the left and right putamen coactivations in the domains of language and execution of speech also revealed that the left putamen exhibited a primarily left-lateralized coactivation network spanning regions associated with language processing (Viñas-Guasch & Wu, 2017). The authors implicated the left putamen particularly with semantic processes. The left caudate nucleus has also been associated with language processing, particularly learning a second language (Tan et al., 2011) and control processes in bilingualism such as code switching (Crinion et al., 2006; Zou, Ding, Abutalebi, Shu, & Peng, 2012), and lesions in the left caudate nucleus were shown to correlate with speech and language impairments following stroke, implicating this region in control processes involving speech and language (Grönholm, Roll, Horne, Sundgren, & Lindgren, 2016). In support of these associations, a DTI study identified a structural network involving the basal ganglia, the thalamus, and Broca’s area, which is argued to support language processing (Ford et al., 2013). Specifically, the authors propose that this basal ganglia-thalamo-Broca’s area system may facilitate semantic and lexical-phonological processes during word selection. Based on these previous studies and the present findings that LBA44 coactivated with these subcortical structures and that LBA44 showed functional specificity for speech, phonology and semantics in the behavioral analysis, it is thought that the basal ganglia-thalamo-LBA44 circuit may support control processes in phonological and semantic processing, and speech.

Although all LIFG ROIs coactivated with parts of the cerebellum, the coactivation peaks for LBA45 and LFo6&7 were parts of a larger cluster in the left hemisphere involving the fusiform gyrus predominantly, but also the left cerebellum (declive) to a much lesser extent, while LBA44 coactivated with the right cerebellum (culmen) specifically. A previous MACM study identified similar coactivation in the culmen overlapping with a more robust cluster in the fusiform gyrus, which was attributed to imprecise allocation of activation in the culmen due to smoothing issues in preprocessing stages especially in adjacent lobes or structures (Bernal et al., 2015). Therefore, the cerebellar coactivation of LBA45 and LFo6&7 in the present study is dubious, whereas LBA44 reliably coactivated with the cerebellum. Indeed, the contrast analysis revealed that only LBA44 specifically coactivated with the right cerebellum. The cerebellum has long been considered to underlie coordination of motor control, while particularly the right cerebellum has recently been associated with mediation of cognitive functions including language (Murdoch, 2010). Neuroimaging research implicated the right cerebellum in speech and language processing (Booth, Wood, Lu, Houk, & Bitan, 2007; Wildgruber, Ackermann, & Grodd, 2001). Furthermore, children with specific language impairment and children with autism spectrum disorder accompanied by language impairment were shown to have morphological differences in the cerebellum compared to their typically developing peers (Hodge et al., 2010). Moreover, a previous r-s fMRI study revealed that Broca’s and Wernicke’s areas were functionally connected to the basal ganglia, the thalamus and the right cerebellum, which were argued to play a role in language processing (Tomasi & Volkow, 2012). The coactivation of LBA44 with the basal ganglia and the cerebellum is consistent with the DP Model (Ullman, 2001, 2004, 2016), according to which the procedural system comprises the basal ganglia and the cerebellum, which are primarily involved in learning and consolidation of procedural memories, on the one hand, and the left premotor cortex (BA6) and posterior LIFG (BA44), which are claimed to process automatized procedural memories, on the other (Ullman, 2016). Specifically, some versions of the DP Model associates the right cerebellum with learning grammatical rules as well as searching lexical items in the declarative system (Ullman, 2001, 2004). Given that recent neuroscience research pointed to the importance of cerebello-basal ganglia-thalamo-cortical loop for motor and nonmotor functions (Bostan, Dum, & Strick, 2013; Bostan & Strick, 2010, 2018; Caligiore et al., 2017; Tomasi & Volkow, 2012), it is thought that the cerebellum may contribute to the basal ganglia-thalamo-LBA44 circuit discussed in the previous paragraph and may mediate and modulate cognitive functions including language (Murdoch, 2010).

### Coactivation patterns of right-hemispheric ROIs

It was found that significantly more papers and experiments were identified for each left-hemispheric ROI compared to the right-hemispheric ones. This is compatible with left-hemispheric dominance for language processing. Also, unlike LIFG ROIs, which revealed a mostly ipsilateral, left-lateralized coactivation network, RIFG ROIs coactivated mostly with contralateral, left-hemispheric regions. This suggests that while LIFG engages an intra-hemispheric language network, RIFG involves an interhemispheric language circuitry, in line with previous investigations (Vigneau et al., 2011, 2006). In parallel with the LIFG results, among RIFG ROIs, RBA44 revealed the most widespread coactivation pattern, followed by RBA45 and, lastly, by RFo6&7. This finding suggests that there is functional segregation within RIFG as part of the language network similar to that of LIFG. Unlike LIFG ROIs, however, there was almost no overlap among all three ROIs in either the left or the right hemisphere, while RBA44 and RBA45 exhibited coactivation overlap in the left and right frontal and insular cortices. This lack of overlap in the coactivation network of all three RIFG subdivisions may constitute one of the functional characteristics that set this region apart from its left-hemispheric homologue and may reflect less consistent involvement in the literature of RIFG in language processing. Nevertheless, RIFG showed a coactivation pattern, somewhat similar to LIFG, within left frontal (IFG, MFG, precentral gyrus, medial frontal regions), insular, parietal (IPL) and temporal (MTG, fusiform gyrus) cortices, as well as subcortical regions (thalamus). This implies that the right hemisphere, particularly RIFG, takes part not only in contextual, holistic and prosodic aspects of language processing as commonly argued in the literature (Dara et al., 2014; George et al., 1996; Lundgren & Brownell, 2016; Parola et al., 2016; Patel et al., 2018; Stockbridge et al., 2021; Xu et al., 2005), but also in lower-level semantic, phonological, syntactic and speech-related processes in conjunction with the left hemisphere. However, given that the behavioral analysis did not reveal any functional specificity for language subdomains within RIFG ROIs, that a disproportionately lower number of language-related studies were identified for RIFG ROIs than LIFG ROIs, and that both LIFG and RIFG ROIs coactivated mostly with left-hemispheric regions, the role of the right hemisphere in language processing seems to be rather limited. In parallel with previous observations (Vigneau et al., 2011), it could be argued that the function of the right cerebral hemisphere within the language network is not language-specific, but may relate to executive processes that are recruited in a task-dependent manner. Along similar lines, an fMRI study revealed that activation in several bilateral regions including IFG, MFG, the precentral gyrus and the insula increased as familiarity of the sentences read decreased (Lai, Van Dam, Conant, Binder, & Desai, 2015). The brain regions bilaterally associated in that study largely overlaps with the right and the left frontal regions coactivating with the RIFG ROIs in the present investigation. The authors of that study associated these right hemispheric regions particularly including right frontal and insular regions with increased cognitive demands of processing unfamiliar stimuli (Lai et al., 2015). Thus, when the current findings are interpreted in light of the previous research, it is thought that RIFG may support mainly left-lateralized language processes when additional cognitive resources are needed.

### Limitations

Several potential limitations pertain to the present investigation and need addressing. First, the BrainMap database, from which the experiments coactivating with IFG ROIs were sampled in this study, represents only a subset of neuroimaging experiments on language processing. This is partially due to the criteria of inclusion in the database. To be eligible for inclusion in the database, published fMRI or PET experiments must report whole-brain coordinates in standard space (MNI or Talairach), which means that experiments reporting only region-of-interest or volume-of-interest results are excluded since they violate the assumption of random spatial distribution across the whole brain (Eickhoff et al., 2012; Müller et al., 2018, 2017). Nevertheless, BrainMap has been expanding continuously with addition of more neuroimaging research by the database staff and by independent researchers into the database, which ensures that it is an extensive and representative sample of neuroimaging studies. Indeed, at the time of the analysis, the functional database of BrainMap comprised 3406 papers corresponding to 16901 experiments and 76016 subjects. In addition, the number of language-related papers identified in the database and included in the MACM analyses was high for most of the ROIs particularly in the left hemisphere (72, 68 and 29 for LBA44, LBA45 and LFo6&7, respectively). However, the number of papers included in the MACM analyses of the right-hemispheric ROIs was lower (22, 21 and 14 for RBA44, RBA45 and RFo6&7, respectively). Of the right-hemispheric ROIs, only RFo6&7 was associated with a lower number of papers than the minimum (around 17-20 experiments) recommended for ALE meta-analyses to obtain enough power for detection of small effect sizes and to prevent over-influence of individual studies (Eickhoff et al., 2016; Müller et al., 2018). Therefore, the coactivation results concerning RFo6&7 should be interpreted cautiously.

Second, the primary objective of the present research was to reveal functional connectivity of IFG subdivisions for language, without specifically testing or contrasting involvement with particular language components (e.g., syntax, semantics, phonology, speech), tasks (e.g., comprehension, production), or stimulus presentation modality (e.g., visual, auditory), which can influence brain-language associations. Addressing all these additional issues at the same time would be a tremendous initiative and surpass the bounds of a single study. Nevertheless, the present findings have been interpreted in relation to certain language components in light of the behavioral analysis and previous research, though these interpretations should be approached tentatively. Future MACM studies may look into the coactivation patterns of LIFG subdivisions for different linguistic components such as syntax and semantics, which would further elucidate brain-language associations and enable a more specific test of neurocognitive models of language processing from a network perspective.

## Conclusion

Utilizing the BrainMap functional database of neuroimaging experiments and meta-analytic connectivity modeling, the present investigation aimed to elucidate functional connectivity profiles of LIFG and RIFG subdivisions (pars opercularis / BA44, pars triangularis / BA45, and pars orbitalis / Fo6&7) during language tasks. A predominantly left-lateralized coactivation pattern was identified for both left- and right-hemispheric ROIs, underscoring the left-hemispheric dominance for language processing. Specifically, differences were revealed among the functional networks of LIFG subdivisions, with posterior-dorsal LIFG (BA44) coactivating with a more extensive dorsal network of regions, particularly spanning bilateral frontal, bilateral parietal, left temporal, left subcortical (thalamus and putamen), and right cerebellar regions, but anterior-ventral LIFG (BA45 and Fo6&7) showing an exclusively left-lateralized involvement of frontal and temporal regions. The findings highlight the extensive cortical and subcortical functional network underlying language and suggest functional segregation within LIFG, with LBA44 acting as a network hub with diverse cortical, subcortical and cerebellar connections as part of the language network. Overall, the present findings shed light on the functional circuitry of language and allowed scrutiny of the predictions made by neurocognitive models of language processing. Also, the functional circuitry of language identified here for healthy participants may also serve as a baseline against which the language network in clinical populations can be compared, potentially facilitating assessment of clinical outcomes in neurorehabilitation of language disorders such as aphasia.

## Footnotes

1 Please see the color bars in Figure 1 for the minimum and maximum probabilities. The mean probabilities (SDs in parentheses) and sizes of the ROIs were as follows: LBA44: 0.77 (0.09), 2409mm^3^; LBA45: 0.64 (0.12), 2363mm^3^; LFo6&7: 0.68 (0.12), 2353mm^3^; RBA44: 0.72 (0.11), 2391mm^3^; RBA45: 0.77 (0.09), 2310mm^3^; RFo6&7: 0.64 (0.11), 2372mm^3^.

